# Linking minimal and detailed models of CA1 microcircuits reveals how theta rhythms emerge and how their frequencies are controlled

**DOI:** 10.1101/2020.07.28.225557

**Authors:** Alexandra P Chatzikalymniou, Melisa Gumus, Anton R Lunyov, Scott Rich, Jeremie Lefebvre, Frances K Skinner

**Affiliations:** Krembil Research Institute, University Health Network; Department of Physiology, University of Toronto; Departments of Medicine (Neurology) and Physiology, University of Toronto

## Abstract

The wide variety of cell types and their inherent biophysical complexities pose a challenge to our understanding of oscillatory activities produced by cellular-based computational models. This challenge stems from the high-dimensional and multi-parametric nature of these systems. To overcome this issue, we implement systematic comparisons of minimal and detailed models of CA1 microcircuits that generate intra-hippocampal theta rhythms (3-12 Hz). We leverage insights from minimal models to guide detailed model explorations and obtain a cellular perspective of theta generation. Our findings distinguish the pyramidal cells as the theta rhythm initiators and reveal that their activity is regularized by the inhibitory cell populations, supporting an ‘inhibition-based tuning’ mechanism. We find a strong correlation between the pyramidal cell input current and the resulting LFP theta frequency, establishing that the intrinsic pyramidal cell properties underpin network frequency characteristics. This work provides a cellular-based foundation from which *in vivo* theta activities can be explored.

## Introduction

Hippocampal theta rhythms (≈ 3-12 Hz) as observed in local field potential (LFP) recordings are associated with cognitive processes of memory formation and spatial navigation (***Colgin, 2013***, ***2016***; ***Hinman et al., 2018***). However, exactly how theta rhythms emerge is a complicated and multi-layered problem. The medial septum (MS) is believed to act as a pacemaker since theta rhythms in the hippocampus are severely attenuated when the MS is lesioned (***Winson, 1978***). Moreover, the various cell types in the MS and in the hippocampus are interconnected in cell-specific ways (***Chamberland et al., 2010***; ***Huh et al., 2010***). This underlines the importance of considering how cellular specifics contribute to theta rhythm circuit dynamics and ultimately function, especially since sophisticated experimental techniques continue to uncover the diversity and distinctness of neurons (***Harris et al., 2018***; ***Hodge et al., 2019***; ***Kepecs and Fishell, 2014***; ***Sugino et al., 2019***).

It is now well-documented that theta rhythms can be generated intra-hippocampally, emerging spontaneously from an isolated whole hippocampus preparation *in vitro* (***Goutagny et al., 2009***). Two computational modelling studies have captured these intrinsic theta rhythms. The first study by ***Ferguson et al. (2017)*** used minimal network models of biophysically simplified neurons, while the second study by ***Bezaire et al. (2016b)*** used biophysically detailed network models. These models can help us understand how these rhythms are generated while taking into consideration each model’s advantages and challenges.

The minimal model of ***Ferguson et al. (2017)*** represents a ‘piece’ of the CA1 region of the hippocampus, and it was developed and constrained against data from the whole hippocampus preparation (***Ferguson et al., 2013, 2015b***). We used this model to examine what ‘building block’ features could underlie theta rhythms (***Ferguson et al., 2015a***, ***2017***). It was found that spike frequency adaptation (*SFA*) and post-inhibitory rebound (*PIR*) building block features of excitatory, pyramidal (PYR) cells in large minimally connected recurrent networks with fast-firing, parvalbumin-expressing (PV+) inhibitory cells could produce theta frequency population rhythms. Specifically, for the model to be consistent with experimental observations of excitatory postsynaptic current (EPSC) and inhibitory postsynaptic current (IPSC) amplitude ratios, the connection probability from PV+ to PYR cells is required to be larger than from PYR to PV+ cells. The minimal model design, strategy and setup suggests that the theta oscillation generation mechanism could be due to *SFA* and *PIR* building block features. However, the challenge is to determine how these insights could apply in the biological, hippocampal system with its larger complement of diverse inhibitory cell types and additional biological details.

The detailed model of ***Bezaire et al. (2016b)*** is a full-scale biological model of the CA1 hippocampus with 338,740 cells that includes PYR cells, PV+ basket cells (BCs), axo-axonic cells (AACs), bistratified cells (BiCs), cholecystokinin-expressing (CCK+) BCs, Schaeffer Collateral-associated (SCA) cells, oriens-lacunosum-moleculare (OLM) cells, neurogliaform (NGF) cells, and ivy cells. The model provides a realistic representation of the hippocampus which is grounded upon a previously compiled, extensive quantitative analysis (***Bezaire and Soltesz, 2013***). It describes the activities of the PYR cells and the eight inhibitory cell types during theta rhythms. In broad terms, this model distinguishes the importance of certain cell types against others, and predicts that cell type variability is necessary for theta rhythms to occur. However, the very complexity of the detailed model poses a challenge in the deciphering of the exact mechanism of the theta rhythm it produces.

The goal of the present paper is to combine the advantages of minimal and detailed models to obtain a cellular-based understanding of theta rhythm generation in the biological system. The strategy we employ is schematized in ***Figure 1***, and the pipeline flow of the paper can be illustrated by three main steps. We first extend the minimal model, **step 1**, to test the robustness of the theta rhythms in the face of PYR cell heterogeneity. This allows us to propose an ‘inhibition-based tuning’ mechanism that underlies theta rhythm generation and frequency control. We next compare minimal and detailed models, **step 2**, to identify commonalities and differences in their structure. Finally, in **step 3**, we extract a ‘piece’ of the detailed model to create the segment model which is comparable in cell numbers to the minimal model, and we investigate the effect of the noted differences on theta. Following a principled exploration of the segment model, we decipher how its theta rhythm is produced. We reveal a strong correlation between the PYR cell net input current and the frequency of the resulting theta rhythm and show that the initial spark of the theta LFP rhythm is due to the PYR cell networks. The inhibitory cell populations on the other hand ‘regularize’ the theta rhythms and increase their power. Not surprisingly, we find degeneracy in our segment models but comparisons with additional experimental observations support some model parameter sets and not others.

**Figure 1.**
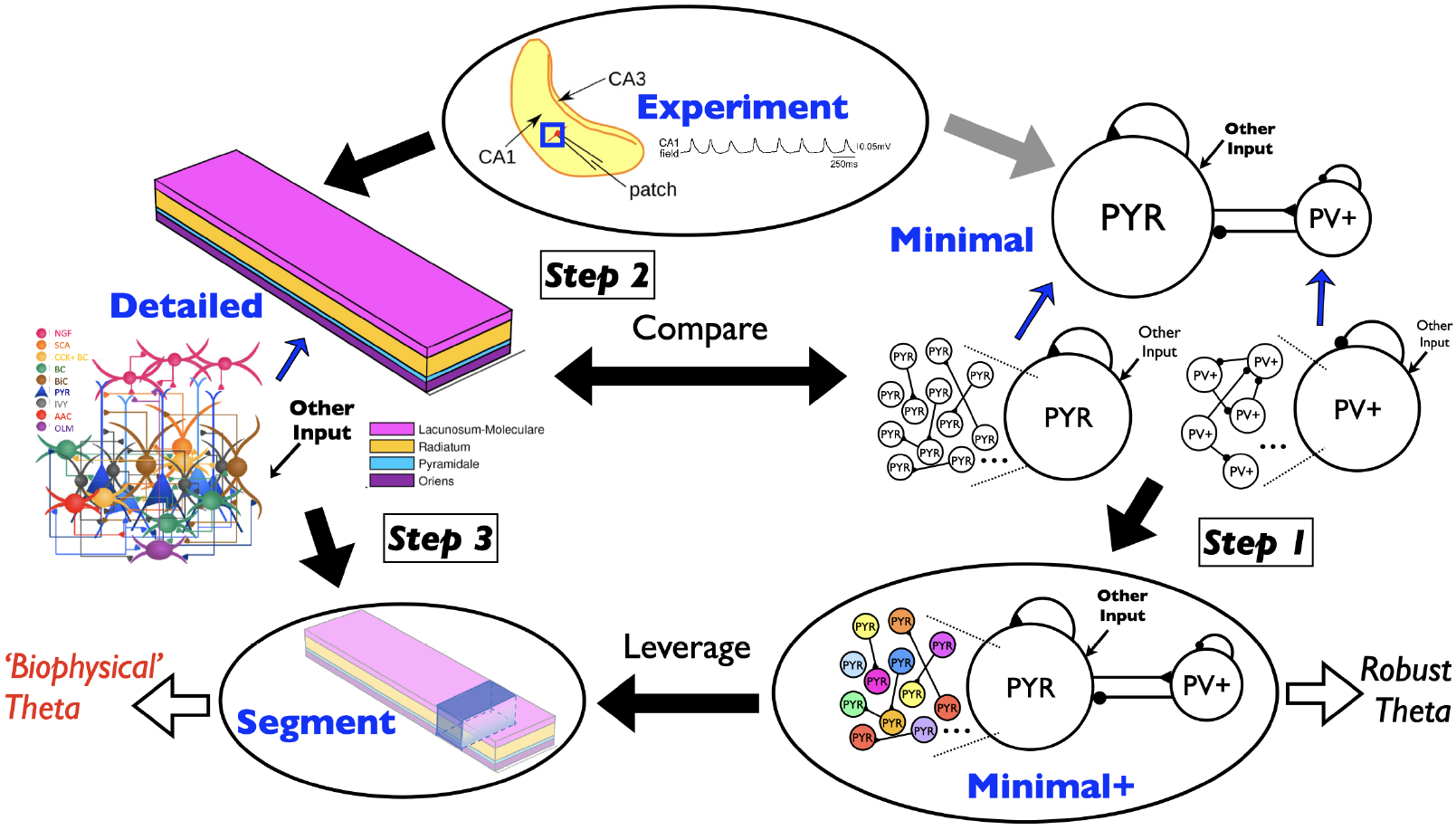
Schematic illustrating overall paper flow and strategy. The experimental context and four model types are referred to in the paper: *Experiment* - a whole hippocampus preparation that exhibits spontaneous theta rhythms (***Goutagny et al., 2009***); *Minimal* - a previously published work of minimal network models representing a ‘piece’ of the whole hippocampus (blue square in *experiment* illustration) that generates theta rhythms within experimental constraints (***Ferguson et al., 2017***); *Minimal+* - an expansion of the minimal model using heterogeneous PYR cells (as illustrated with differently coloured PYR cells) that is used in the present paper; *Detailed* - a previously published work of a full-scale detailed CA1 microcircuit model (eight different inhibitory cell types and PYR cells) that generates theta rhythms without any oscillatory input (***Bezaire et al., 2016b***); and *Segment* - a network model representing a ‘piece’ of the detailed model, that is used in the present paper. The three main steps in the flow of the paper are shown (**Steps 1-3**), and the foci of the work in the present paper are illustrated by the black arrows: The detailed model is examined in light of the experimental data;a systematic comparison between minimal and detailed models is done; the segment model is created from the detailed model; the minimal+ model is constructed based on the minimal model, and mechanistic insights resulting from the minimal+ model are leveraged in the segment model. The black open arrows illustrate that ‘Robust Theta’ in the minimal+ model is examined leading to hypothesis development, and leveraging this in the segment model helps with an understanding of ‘Biophysical Theta’ where multiple cell types can be considered. The grey arrow illustrates previously done work where the minimal model was developed and examined in light of the experimental data (***Ferguson et al., 2017***). Illustrations include: *Minimal* model setup with PYR and fast-firing PV+ cells, *Detailed* model setup with 9 cell types (NGF, SCA, CCK+ BC, BC, BiC, PYR, IVY, AAC, OLM) and layer-specific connectivity, *Experiment* of whole hippocampus preparation with a LFP theta example, heterogeneous PYR cells as different colors in *Minimal+* model, and a shaded portion of the *Detailed* model prism to illustrate the *Segment* model. Acronyms are defined in the main text. This figure is adapted from parts of other figures: Figs. 1 & 8 of ***Huh et al. (2016)***, Fig. 2 of ***Ferguson et al. (2017)***, and Fig. 1 of ***Bezaire et al. (2016b)***.

Overall, we have combined minimal and detailed models to establish a cellular basis for how the theta rhythms could be robustly generated and how their frequency is controlled in the biological system. By extension we have identified common principles of the theta generation mechanism between the two models and we discuss their differences. Moving forward, this work provides a solid biological ‘seed’ from which to examine the multi-layered aspects of theta rhythms in the hippocampus.

## Results

The flow of the results section is as follows. We begin by exploring the robustness of the theta rhythm in the minimal model from the perspective of its building block features. Subsequently, phase response curve (PRCs) analysis leads to the proposition of an ‘inhibition-based tuning’ mechanism of theta rhythm generation and frequency control. To investigate this mechanism in the detailed model, we do the following. First, we compare EPSC/IPSC amplitude ratios in the detailed model with those in the whole hippocampus preparation as it was already done with the minimal model. Next, we carry out a systematic comparison between minimal and detailed models by comparing connectivities, synaptic weights and external drives. Finally, we isolate a ‘piece’ of the detailed model - the segment model - comparable in cell numbers to the minimal model. We examine the segment model in a principled manner according to minimal and detailed model comparisons. As the segment model is much smaller than the detailed model, we can perform extensive explorations and establish how intra-hippocampal theta rhythms are generated and how their frequencies are controlled.

### Robustness of theta generation in the minimal model

The minimal model suggested that the generation of theta oscillations could be based on the amount of spike frequency adapation (*SFA*) present in the pyramidal (PYR) cells together with their ability to exhibit post-inhibitory rebound (*PIR*) in large networks of minimally connected PYR cells, interconnected with parvalbumin positive (PV+) fast-firing inhibitory cells (***Ferguson et al., 2017***). Inherent with *SFA* and *PIR* building block features is a rheobase (*Rheo*) feature, which is the amount of current required to make the PYR cell spike (derived from fitting to the experimental data in (***Ferguson et al., 2015b***)). However, in this previous study we did not specifically examine the sensitivity of theta rhythms to these building block features (*SFA, Rheo, PIR*).

The minimal model used an Izhikevich mathematical model structure for the cellular representations (***Izhikevich, 2006***), and while it did not have any direct biophysical ion channel equivalents, its frequency-current (f-I) curve was fit to electrophysiological recordings of PYR cells in the the whole hippocampus preparation (***Ferguson et al., 2015b***). The PYR cell model parameter values, herein referred to as default values, are: *a*=0.0012; *b*=3.0, *d*=10, *k_low_*=0.10. We used a straightforward approach to quantify the *SFA, Rheo, PIR* building block features (see Methods). For the PYR cell model with default parameter values, the quantified building block feature values are: *SFA*= 0.46 Hz/pA; *Rheo*= 4.0 pA;*PIR* = −5.0 pA. We refer to these values as *base* building block feature values. The larger the quantified *SFA* value is, the stronger is the amount of the PYR cell adaptation, i.e., we get more reduction in the PYR cell spike frequency for a fixed amount of input current. The more negative the quantified *PIR* value is, the larger is the hyperpolarizing step required to generate a spike at the end of the step.

#### Examining the contribution of building block features

In the extensive network simulations of ***Ferguson et al. (2017)***, the PYR cell models were homogeneous in terms of their (*a, b, d, k_low_*) model parameter values. However, the networks were not homogeneous because of the noisy external drives to the PYR cell models. Because of its direct connection to the experimental data, the minimal model with its building block features was considered to encompass key ‘biological balances’ important for theta rhythm generation. To examine the robustness of the theta-generating mechanism in the minimal models with consideration of the *SFA, Rheo* and *PIR* building block features, we create heterogeneous PYR cell populations from a model database that is generated by ranging *a, b, d, k_low_* parameter values around default ones. In turn, this model database provides a distribution of quantified *SFA, Rheo, PIR* building block feature values. The distributions of values are shown in ***Figure 2***, and the locations of the base values are indicated by vertical black arrows.

**Figure 2.**
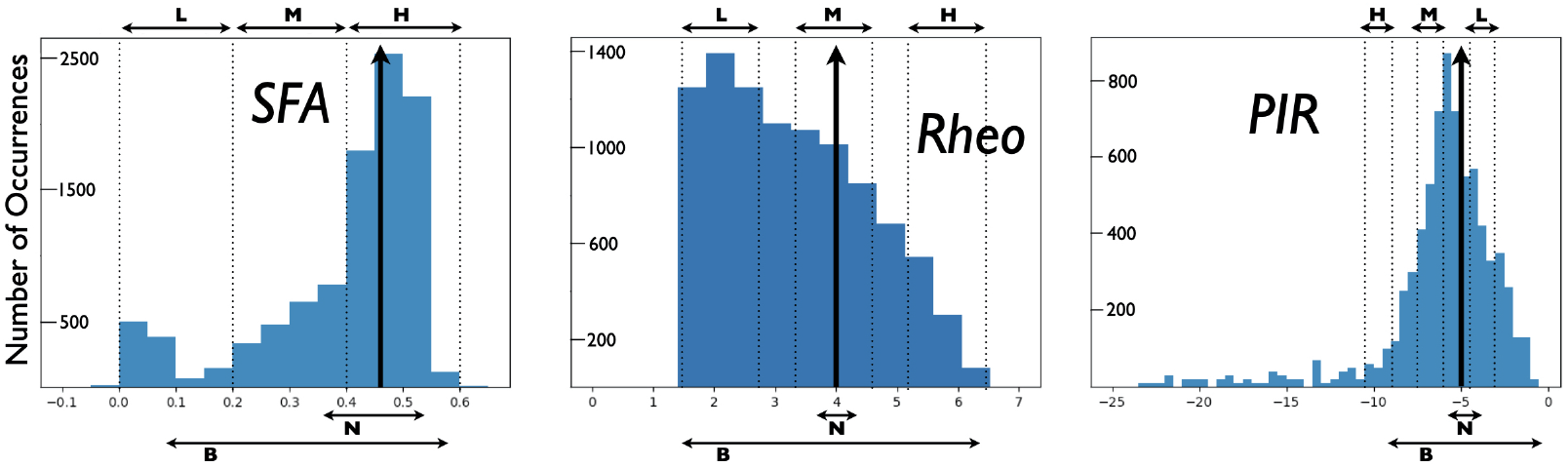
Distributions of PYR cell building block features from created model database. A heterogeneous set of PYR cells was created and their ‘building block’ features of *SFA*, *Rheo* and *PIR* were quantified. Histograms show the number of occurrences of *SFA [=] Hz/pA, Rheo [=]pA, PIR[=] pA* values. See Methods for details of quantifications. Also shown are narrow (*N*) and broad (*B*) subsets used to consider heterogeneous PYR cell populations in one way (i), and low (*L*), medium (*M*), high (*H*) subsets to consider heterogeneous PYR cell populations in another way (ii). See main text for further details. Vertical black arrows indicate [*SFA,Rheo,PIR] base values* of a PYR cell model with default model parameters. *SFA* histogram has a bin resolution of 0.05, and *Rheo, PIR* histograms have a bin resolution of 0.5. Acronyms are defined in the main text.

Before delving into heterogeneous excitatory-inhibitory (E-I) model networks, let us first examine E-I networks of homogeneous PYR cell models with parameter values different from the default ones, but with similar values for quantified building block features. The resulting networks produce clear population bursts, but with some variation in frequency and power. Specific examples are shown in ***Table 1*** along with their model parameter and quantified building block feature values. The fact that the rhythm is not lost in any of these networks with homogeneous model parameter values already suggests that the populations bursts are not particularly sensitive to the specific *SFA* building block quantified values as the rhythm isn’t lost as *SFA* varies. However *SFA* has some effect on the specific power and frequency of the population bursts.

**Table 1.** E-I Network Simulation Examples with Homogeneous PYR cell models.

| **Homogeneous cells in network**<br>Model ID # | **Parameter values**<br>(*a*, *b*, *d*, *k<sub>low</sub>*)<br>Units: (1/ms, nS, pA, nS/mV) | **Quantified values**<br>(*SFA*, *Rheo*, *PIR*)<br>Units: (Hz/pA, pA, pA) | **Power**<br>(mV<sup>2</sup>/Hz) | **Frequency**<br>(Hz) |
| --- | --- | --- | --- | --- |
| Original (base) | (0.0012, 3.0, 10, 0.10) | (0.46, 4.0, -5.0) | 0.36 | 12.2 |
| # 7 | (0.00072, 3.6, 18, 0.16) | (0.51, 4.0, -5.0) | 0.21 | 11.8 |
| # 32 | (0.00072, 4.8, 12, 0.16) | (0.51, 4.0, -5.0) | 0.37 | 14.2 |
| # 56 | (0.00096, 3.6, 4, 0.12) | (0.38, 4.0, -5.0) | 0.40 | 13.6 |
| # 81 | (0.00096, 4.2, 12, 0.10) | (0.49, 4.0, -5.0) | 0.42 | 13.8 |
| # 115 | (0.0012, 3.6, 14, 0.06) | (0.49, 4.0, -5.0) | 0.34 | 13.0 |

Let us now consider E-I networks with heterogeneous PYR cell models (*Minimal+* models as illustrated in ***Figure 1***). We classify the PYR cells from the created model database in two groups according to their quantified values of the [*SFA, Rheo, PIR]* building block feature trio. The first group corresponds to: (i) Narrow (*N*) or broad (*B*) ranges of [*SFA, Rheo, PIR]* values that include the base values, and the second group corresponds to: (ii) Low (*L*), medium (*M*) or high (*H*) ranges of [*SFA, Rheo, PIR]* values that do not necessarily include the base values. These groups are shown in ***Figure 2***. For each group we create networks corresponding to combinations of the quantified values of the *SFA, Rheo, PIR* building block feature ranges. For (i), there are eight possible E-I network cases from *N* and *B* building block combination sets and the number of models in each case is given in ***Table 2***, along with the frequency and power of the particular network. For (ii), there are 27 possible network cases from *L, M* and *H* building block combination sets and the number of models in each case is also given in ***Table 2***, along with the frequency and power of the particular network. As it turns out, there are no PYR cell models in the created model database for *HHH, HHL, MHH, MHL, LHH, LHL* network cases. We thus have simulation output for only 21 different E-I networks with heterogeneous PYR cell populations generated using (ii). Further details on the model database are given in the Methods.

**Table 2.** Heterogeneous E-I Network Simulations.

| **Network case**<br>*[SFA,Rheo,PIR]* | **Number of**<br>**different**<br>**PYR cell**<br>**models** | **Power**<br>(mV<sup>2</sup>/Hz) | **Frequency**<br>(Hz) |
| --- | --- | --- | --- |
| **Group (i)** |  |  |  |
| *NNN* | 137 | 0.37 | 13.0 |
| *BBB* | 6780 | 0.27 | 13.0 |
| *BBN* | 550 | 0.28 | 12.8 |
| *BNB* | 1010 | 0.29 | 13.0 |
| *BNN* | 180 | 0.34 | 13.4 |
| *NBB* | 4955 | 0.30 | 13.0 |
| *NBN* | 416 | 0.24 | 12.2 |
| *NNB* | 729 | 0.33 | 12.6 |
| **Group (ii)** |  |  |  |
| ***HML*** (R) | 556 | 0.38 | 13.0 |
| *HHM* | 313 | 0.40 | 15.6 |
| *HMM* | 493 | 0.37 | 12.8 |
| *MHM* | 157 | 0.46 | 15.8 |
| ***MMH*** (R) | 25 | 0.19 | 9.6 |
| *MMM* | 294 | 0.31 | 13.2 |
| *MML* (R) | 110 | 0.37 | 13.8 |
| *MLL* * | 99 | 0.12 | 10.0 |
| *LHM* | 49 | 0.35 | 16.2 |
| *LMH* * | 12 | 0.15 | 9.8 |
| *LMM* | 103 | 0.30 | 13.6 |
| ***LML*** | 74 | 0.32 | 15.0 |
| *LLM* * | 29 | 0.15 | 10.4 |
| *LLL* | 64 | 0.17 | 12.0 |
| **No Rhythm** |  |  |  |
| *HMH* (R-supp) | 33 | 0.06 | n/a (9.2) |
| *HLH* | 97 | 0.01 | n/a (1.2) |
| *HLM* (R) | 171 | 0.01 | n/a (0.6) |
| *HLL* | 417 | 0.02 | n/a (0.6) |
| *MLH* (R-supp) | 27 | 0.04 | n/a (8.2) |
| *MLM* (R-supp) | 50 | 0.07 | n/a (8.6) |
| *LLH* (R-supp) | 16 | 0.08 | n/a (10.0) |

There is a clear maintenance of rhythms for the eight cases of heterogeneous group (i), as shown in the top part of ***Table 2***, where the building block quantified values are chosen in either a narrow or broad fashion encompassing base values. Their frequencies are similar to each other and to that of the E-I network with homogeneous, default PYR cell model parameter values (see first row in ***Table 1***). Interestingly, the network power is larger when there is a narrow rather than a broad range of values encompassing base values (compare *NNN* to *BBB* in ***Table 2***), suggesting that particular quantified building block feature values are important for the presence of robust theta frequency population bursts. In essence, these simulations indicate that the theta generation mechanism in the minimal model is robust. That is, if we have heterogeneous E-I model networks with PYR cell model parameter values that have broadly distributed building block feature values that include base values, then population rhythms remain with less than a 1 Hz variation in population frequency. This further implies that a quantification of the building block features can capture the underlying E-I balances necessary for the emergence of theta frequency population bursts in the minimal model.

Let us now examine the output for the 21 cases of heterogeneous E-I networks with PYR cell models that have quantified building block feature values that do not necessarily encompass base values, i.e., group (ii). This is shown in ***Table 2*** where it is clear that a rhythm (i.e., population bursts) is not always present. We first note that the E-I network for the *HML* case is the one that mostly encompasses base values for all three building block features. As one might expect, the power and frequency of this E-I network case is similar to the heterogeneous (i) E-I network cases which also encompass the base values. Considering the network power values of all of these heterogeneous network cases, it is easy to see which networks are not rhythmic. Essentially, if the power is below 0.1, then there is not a clear rhythm - these cases are shown in the lower part of ***Table 2***. The cases in ***Table 2*** that are starred are networks that have started to lose their rhythm. To view the output from several heterogeneous E-I networks, in ***Figure 3*** we show PYR cell raster plots for four cases (designated with an ‘R’ in ***Table 2***). In three of them, there is still a rhythm, but there are clear frequency and PYR cell burst firing characteristic differences. In ***Figure 3-Figure Supplement 1*** we show PYR cell raster plots for four additional cases (designated with an ‘R-supp’ in ***Table 2***) for when the rhythm is lost so that the different patterning can be seen.

**Figure 3.**
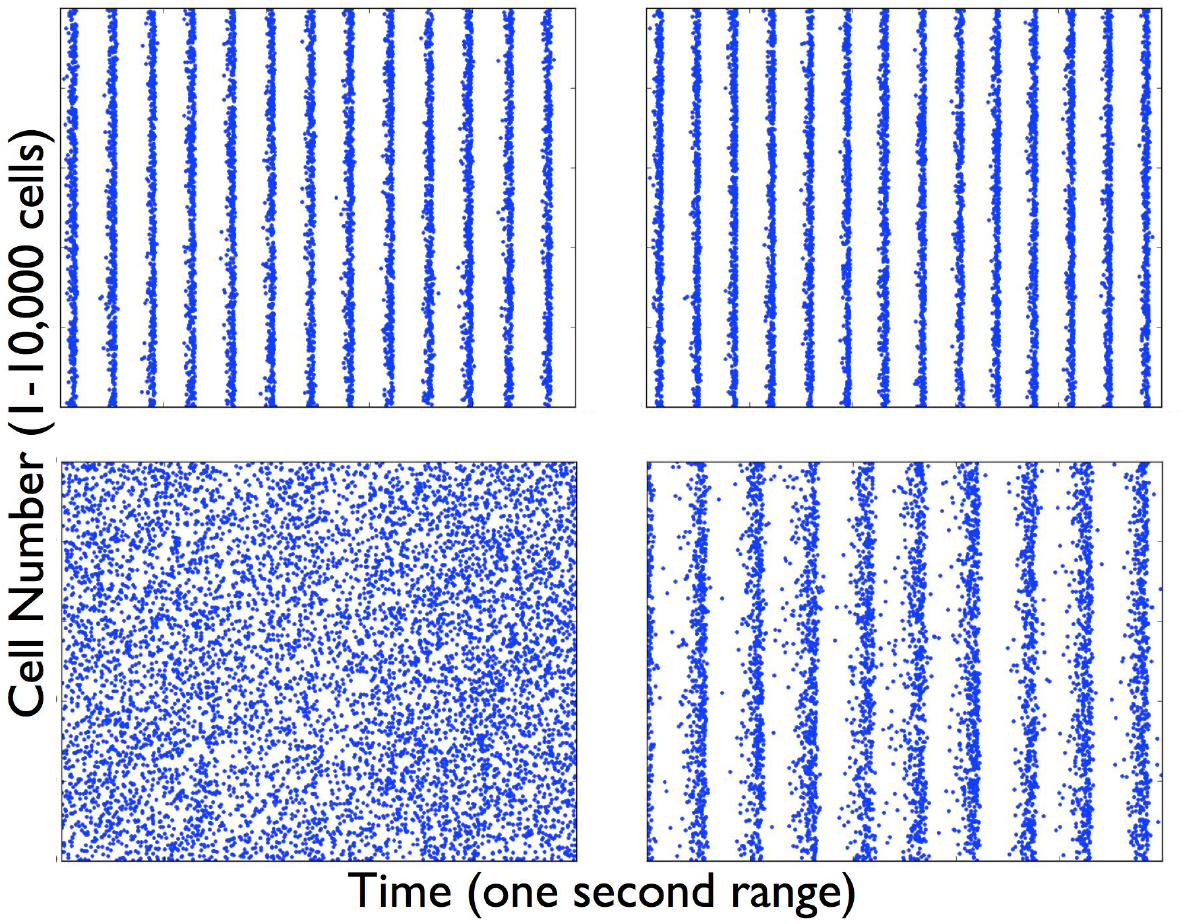
Raster plots of PYR cells in heterogeneous E-I networks. Simulations of E-I networks with 10,000 heterogeneous PYR cells and 500 PV+ cells produce output that have PYR cell raster plots as shown here with a one second time range. The specific examples are labelled as (*R*) in ***Table 2*** and refer to the following sets: HML (top-left), MML (top-right), HLM (bottom-left), MMH (bottom-right). Acronyms are defined in the main text. **Figure 3-Figure supplement 1**. *Loss of Rhythm - Raster plots of PYR cells in heterogeneous E-I networks*.

In considering the cases in which the rhythm is lost, it appears that the existence of the rhythm is not heavily dependent on the specific *SFA* quantified values, since rhythms still exist even when moving away from “Hxx” cases (i.e., those encompassing the base *SFA* value) - *MML* and *LML* cases. However, the rhythm *is* lost if the E-I networks do not include base values for *Rheo* or *PIR*. Specifically, “xMx” (base *Rheo* value) or “xxL” (closest to base *PIR* value) cases. For *Rheo*, consider the *HLL* case (no *HHL* case to consider) and for *PIR*, consider the *HMH* case (less so for the *HMM* case). This allows us to express the following: the particular rheobase current value of the PYR cell, and the ability of the PYR cell to fire a spike with a less hyperpolarized current step are needed for the theta-generating mechanism in the minimal models, along with some amount of spike frequency adaptation.

In summary, these simulations of E-I networks with heterogeneous PYR cell populations have allowed us to gauge the contributions of the different building block features and have helped us to confirm the robustness of the theta-generating rhythm mechanism. As a result, we can reasonably establish that theta frequency population bursts in the minimal model are particularly sensitive *PIR* and *Rheo* feature values, and less sensitive to *SFA* values. Let us now examine how the frequency of the population rhythm could be controlled.

#### Using PRCs to develop a hypothesis of theta frequency control

We have now determined that specific quantified values for *Rheo* and *PIR* building block features are important for theta population rhythms. The *PIR* building block feature is quantified as the size of a hyperpolarizing current step required to evoke a spike (see Methods). We note that this does not necessarily mean that the PYR cells fire due to inhibitory inputs from the PV+ cells during ongoing theta rhythms. In the Izhikevich cell model structure, the ability of a cell to spike after an inhibitory step is reflected in the *b* parameter (see equations in Methods), which needs to be positive for the PYR cell to fire after a hyperpolarizing step. To examine whether the PYR cells in the network fire due to the inhibitory input they receive, we compare the timing of the PYR cell spikes relative to the timing of their incoming IPSCs. Examples are shown in ***Figure 4*** on two different timescales. From them, we can say that the PYR cell firing does not specifically occur *because* of their IPSCs, as spiking can occur before or just after its IPSCs. Due to the limited nature of the minimal model, it is not helpful to carry out comparisons of EPSC and IPSC values relative to experiment. Even though we had previously found that the EPSC/IPSC amplitude ratios were experimentally appropriate for both PYR and PV+ cells ***Ferguson et al. (2017)***, the limited nature of the minimal model prohibits us from probing exact experimental values of EPSCs and IPSCs. Instead, to get a further understanding on E-I balances dictating the frequency of the theta rhythm, we turn to PRC considerations (***Schultheiss et al., 2011)***.

**Figure 4.**
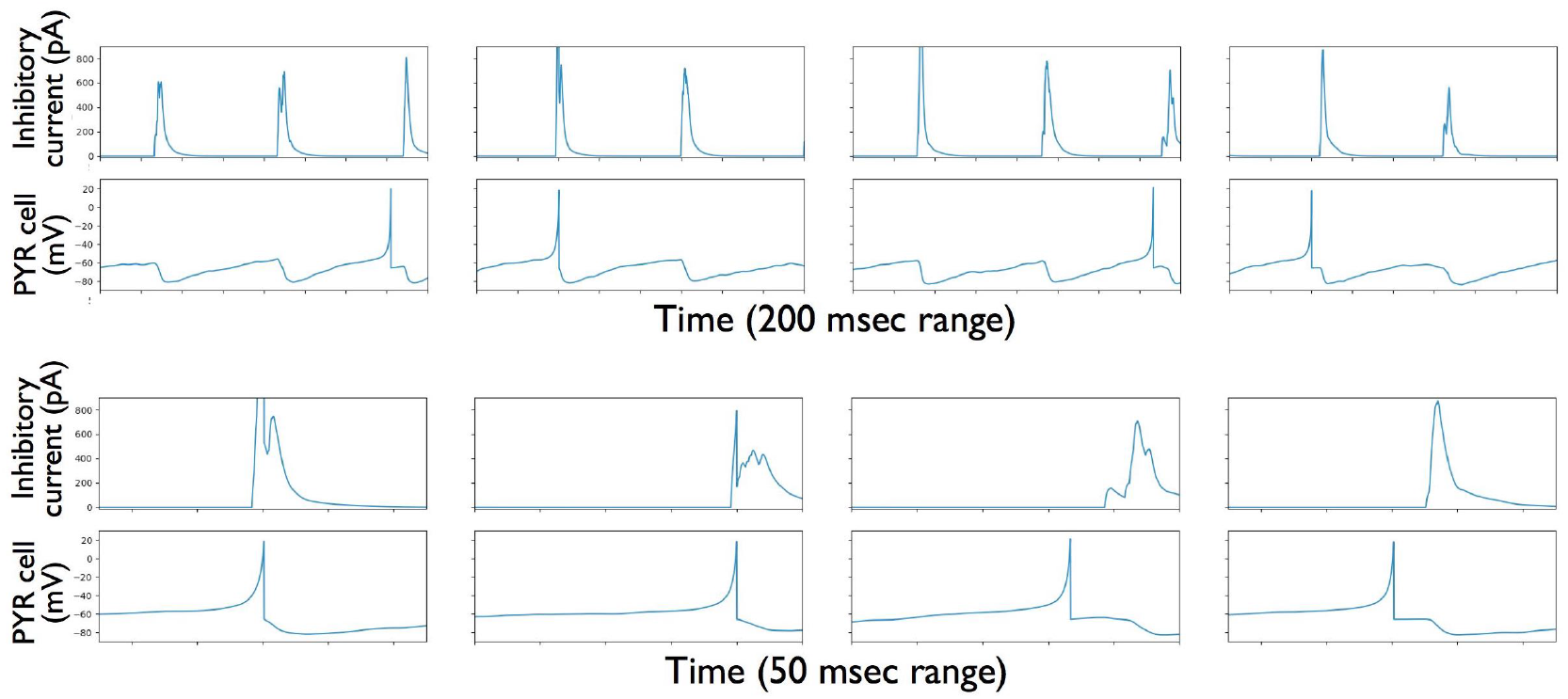
Examples of inhibitory currents onto PYR cells together with PYR cell membrane voltages. Four examples of a PYR cell’s membrane voltage and the inhibitory current (IPSC) onto it. A PYR cell spike can be seen in each example. The top row is shown for a 200 msec time range, and the bottom row is for the same example, but for a 50 msec time range that includes the PYR cell spike. The IPSC can be clearly seen as occurring either after or just before the spike. The PYR cell is one of the 10,000 PYR cells in the heterogeneous E-I network, *BBB* set. Acronyms are defined in the main text.

We hypothesize that the PYR cell network is generating population bursts on its own (given its cellular adaptation characteristics) with the PV+ cell network providing an inhibitory ‘bolus’. We thus consider that the resulting frequency of the E-I network’s population bursts is due to a combination of the PYR cell’s firing frequency combined with how much an inhibitory input could advance or delay the PYR cell spiking. This setup is schematized in ***Figure 5*** as follows: Each PYR cell in the network receives excitatory input from other PYR cells as well as a noisy excitatory drive. The amount of input a PYR cell receives would of course fluctuate over time, but consider that the PYR cell receives a mean excitatory input of about 20 to 30 pA based on parameter values of the minimal models. In these models theta population bursts occur when PYR cells receive a zero mean excitatory drive with fluctuations of ≈ 10-30 pA (***Ferguson et al., 2017***). We generate PRCs by considering an inhibitory ‘bolus’ that a PYR cell would receive by the inhibitory PV+ cell population in the minimal model. The inhibitory pulse would advance or delay the subsequent PYR cell’s spike as given by the PRC. Further details are provided in the Methods.

**Figure 5.**
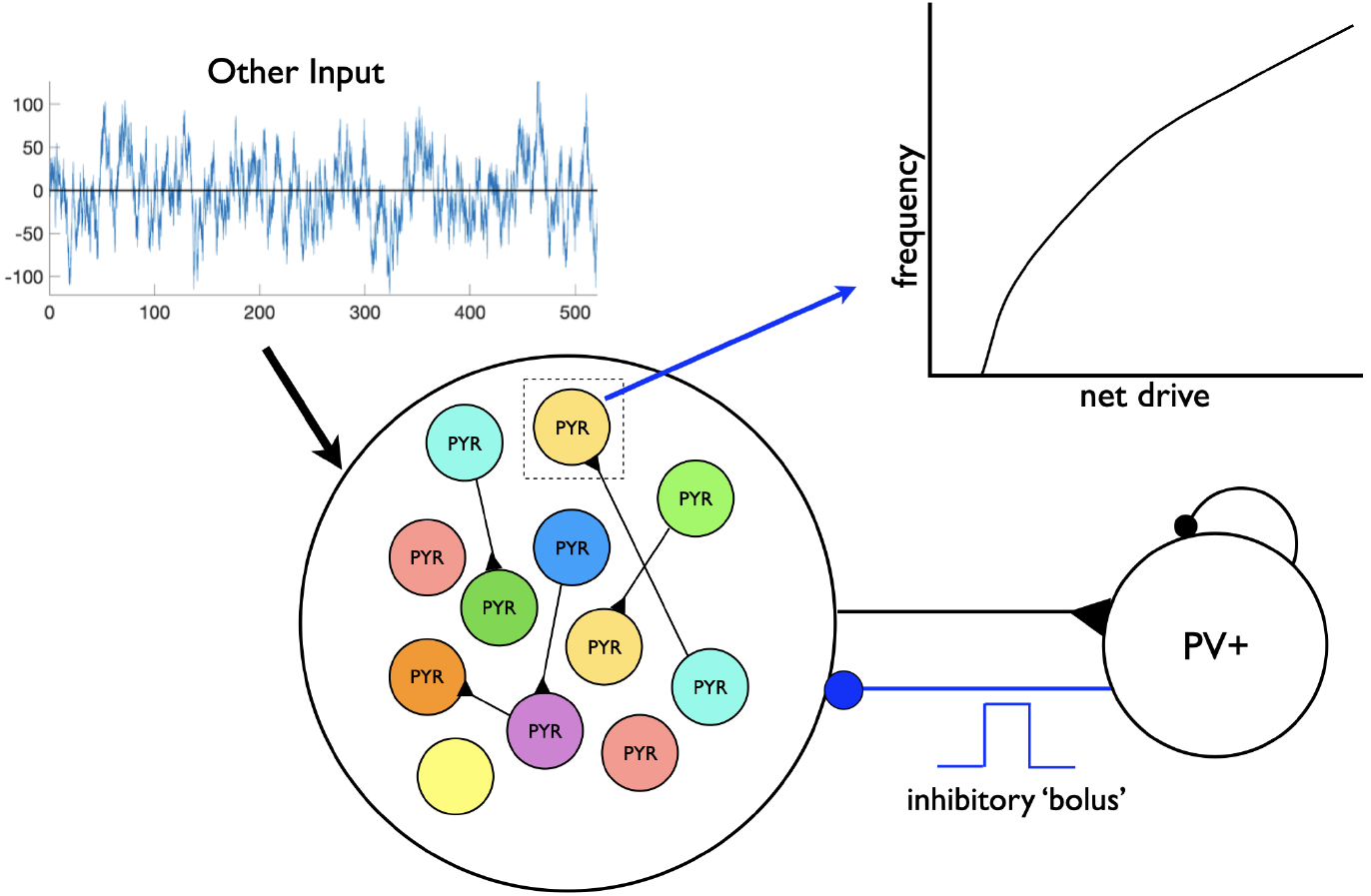
Schematic of setup for phase response curve (PRC) calculations. Using the minimal model structure, and assuming a theta-generating mechanism based on *SFA*, PRCs are generated based on an inhibitory input (‘bolus’) coming from the PV+ cell network to a PYR cell in the PYR cell network. Each PYR cell is receiving a noisy, excitatory drive shown as ‘Other Input’, and an illustrative f-I curve for a PYR cell is shown. The f-I curve with a specified net drive would dictate the result of the computed PRC based on the inhibitory input. Acronyms are defined in the main text.

We consider three cases of heterogeneous E-I networks which exhibit different population burst frequencies. The first case is the *MMH* network with a ‘slow’ frequency of 9.6 Hz, the second case is the *HML* network with a ‘medium’ frequency of 13 Hz, and third case is *LML* network with a ‘fast’ frequency of 15 Hz ***Table 2***. We generate PRCs for the PYR cell models in each of these three cases. Each PYR cell model has particular PRC characteristics due to its *a,b,d,k_low_* model parameter values and exhibits a specific intrinsic frequency for a given input. The calculation of these PRCs is described in the Methods. In ***Figure 6*** we show differences between PRC properties and individual cell firing frequencies for each of the three cases, using an input current of 30 pA. The PRCs for each case show distinct features: for instance, the PYR cells in the *HML* case uniquely exhibit a region of phase-advance, while the PYR cells in the *LML* case have the largest phase delay for perturbations delivered at all but the latest phases. These PRC examinations provide evidence in support of the notion that the frequency of the E-I network population burst is strongly affected by the intrinsic properties of the PYR cells. For instance, while the PYR cells in the *LML* case have the fastest individual firing frequencies (notably faster than what is seen in population models), their PRCs may be slowing down this frequency by means of the inhibitory ‘bolus’ of synaptic inhibition. Meanwhile, the PYR cells in the *HML* case have the slowest individual firing frequencies, although they participate in ‘medium’ speed theta rhythms. The PRC in this case, particularly the region of phase-advance, may play a role in accelerating the PYR cells by means of their inhibitory synaptic input. Frequencies and PRCs for a different input current (20 pA) are shown in ***Figure 6-Figure Supplement 1***.

**Figure 6.**
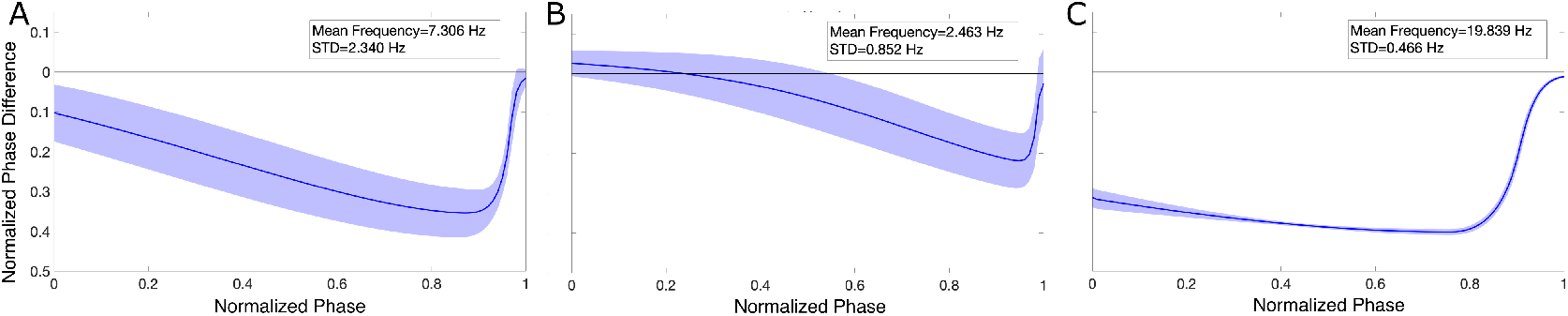
PYR cells from three heterogeneous E-I network cases show distinct PRC features and firing frequencies. Mean PRC (solid line) for PYR cells of a particular case (*MMH* in panel **A**, *HML* in panel **B**, and *LML* in panel **C**) calculated with an input current of 30 pA, with the shading representing ± the standard deviation. The mean and standard deviation of the firing frequencies of the PYR cells at this input level are included in the inset of each panel. PYR cells of the *MMH* case produce ‘slow’ population theta frequency, and PYR cells exhibit moderate individual firing frequencies but notably only show phase-delay in their PRCs. PYR cells of the *HML* case produce ‘medium’ population theta frequency, and PYR cells show the slowest individual firing frequencies, but the region of phase-advance in their PRCs reveals a potential mechanism by which these frequencies might be increased in the network setting. Finally, PYR cells of the *LML* case produce ‘fast’ population theta frequency, and PYR cells show the highest individual firing frequencies, with a potential mechanism by which these are slowed in the network setting revealed by the PRCs with the most marked phase-delay. Acronyms are defined in the main text. **Figure 6-Figure supplement 1**. *PRCs calculated with a 20 pA input show similar features in the three PYR cell populations*.

In essence, this PRC examination allows us to propose that the frequency of the network population bursts depends on the net amount of input delivered to the PYR cells, including the inhibitory bolus. In other words, the frequency response depends on the intrinsic properties of the PYR cells, as given by its f-I curve. This in turn implies that a stable population burst is achieved if the excitation and inhibition are balanced so that proper inhibitory ‘tuning’ can take place. However, just from these minimal model examinations, it is unclear whether such a relationship between PYR cell inputs and network frequency would exist in biologically realistic networks.

Overall, our expansion of the minimal model to include heterogeneous PYR cell populations (see ***Figure 1***) revealed a robustness in the emergence of theta rhythms, and uncovered a sensitivity to the specific quantifed values of the *PIR* and *Rheo* building block features, but not to *SFA*. The use of PRCs showed that the resulting frequency of the population bursts could be largely due to PYR cell intrinsic properties. These explorations in the minimal model lead us to hypothesize an ‘inhibition-based tuning’ mechanism underlying the robust emergence of intrinsic, intra-hippocampal theta rhythms and their frequency control. In this mechanism, together with *SFA, Rheo* and *PIR* building block features, two key aspects are important: (i) The PYR cell population needs to be large enough so that it can collectively generate a strong excitatory drive to the inhibitory PV+ cells. In turn, the PV+ cell population should be able to fire enough (and coherently) to create a strong inhibitory ‘bolus’ that tunes and regularizes the PYR cell population bursting output. (ii) The net input (recurrent excitation, excitatory drive, incoming inhibition) received by the PYR cell situates it in a frequency range that allows theta frequency population bursts to occur. The resulting theta frequency of population bursts are fundamentally ‘controlled’ by the net amount of input that the PYR cells receive.

### Linkage explorations between minimal and detailed models generating intrinsic theta rhythms intra-hippocampally

With a clear sense of how stable theta frequency population bursts are generated in the minimal model, we turn to the detailed model with its empirically-based connections and biophysical cellular specifics. To consider whether the detailed model uses similar theta-generating mechanisms as the minimal model, we examine commonalities and differences between the two models, as illustrated by ‘compare’ in ***Figure 1***. However, we first turn to an examination of EPSC/IPSC amplitude ratios in the detailed model relative to those observed in the whole hippocampus preparation.

#### EPSC/IPSC amplitude ratios in the detailed model are consistent with those in the whole hippocampus preparation

In the minimal model, when we ‘matched’ model EPSC/IPSC amplitude ratios with experiment (***Huh et al., 2016***), we predicted that connection probabilities from PV+ to PYR needed to be larger than those from PYR to PV+ cells (***Ferguson et al., 2017***). The detailed model is experimentally constrained in a bottom up fashion, using cellular data and connectivity information from a plethora of experimental data (***Bezaire and Soltesz, 2013***). Whether the detailed model yields meso-level measurements, such as EPSC/IPSC amplitude ratios that agree with experimental observations from the whole hippocampus preparation, has not been directly assessed. Thus, we here examine whether the detailed model exhibits ratios that ‘match’ those observed in experiments from the whole hippocampus preparation, as was already considered in the minimal model. From the experimental data it is abundantly clear that the EPSC/IPSC amplitude ratios for PYR cells are much less than for PV+ cells. For the detailed model, we consider PV+ cells to represent BCs, BiCs, or combinations of BCs, BiCs and AACs. We choose 15 cells of each type and extract EPSCs and IPSCs at the somata of the different cell types and compute the ratios. We find that regardless of the PV+ cell type or combination considered, it is always the case that the EPSC/IPSC amplitude ratios are consistent with experiment - larger on PV+ cells than on PYR cells - as shown in ***Table 3***. Further details are provided in the Methods.

**Table 3.** EPSC/IPSC Amplitude Ratios from Detailed Model Network Cells.

| PV+ cell type | EPSC/IPSC amplitude ratio<br>(on PYR cell) |
| --- | --- |
| = BC | 4.05 ± 0.86 |
| = BiC | 7.21 ± 1.19 |
| = BC/BiC | 2.95 ± 0.62 |
| = BC/AAC/BiC | 1.78 ± 0.39 |
| = All inhibitory cell types | 1.32 ± 0.24 |
|  | EPSC/IPSC amplitude ratio<br>(on PV+ cell) |
| = BC | 11.71 ± 2.66 |
| = BiC | 34.97 ± 5.28 |

#### Minimal model connectivity prediction validated using detailed model empirical numbers

In the minimal model we predicted that to have EPSC/IPSC amplitude ratios that are consistent with the experimental observations, it is necessary for the connection probability from PV+ to PYR cells to be larger than from PYR to PV+ cells. The connectivities in the detailed model are based on empirical determinations (***Bezaire and Soltesz, 2013***). Thus, if the minimal model is an appropriate representation of the CA1 microcircuitry, its connection probabilities should be in line with those in the detailed model. To consider this, we note two things. First, the minimal model only includes fast-firing PV+ and PYR cells, and second, it uses a random connectivity scheme. Thus, to make comparisons, we consider only PV+ cell types and PYR cells from the detailed model and determine connection probabilities between them using their empirically-based connection schemes. Three inhibitory interneuron cell types in the detailed model can be considered as fast-firing PV+ cell types. These are the BCs, the BiCs and the AACs. Considering only these three inhibitory cell types and the PYR cells, we extracted the number of their post-synaptic connections. This is shown in schematic form in ***Figure 7***. To compare connection probabilities between minimal and detailed models we considered that the fast-firing PV+ cell type in the minimal model could correspond to: (i) only BCs; (ii) only BCs and AACs; (iii) only BCs and BiCs;(iv) BCs, AACs and BiCs. BCs represent the majority of fast-firing PV+ cell types and so they are included in all of the different combinations.

**Figure 7.**
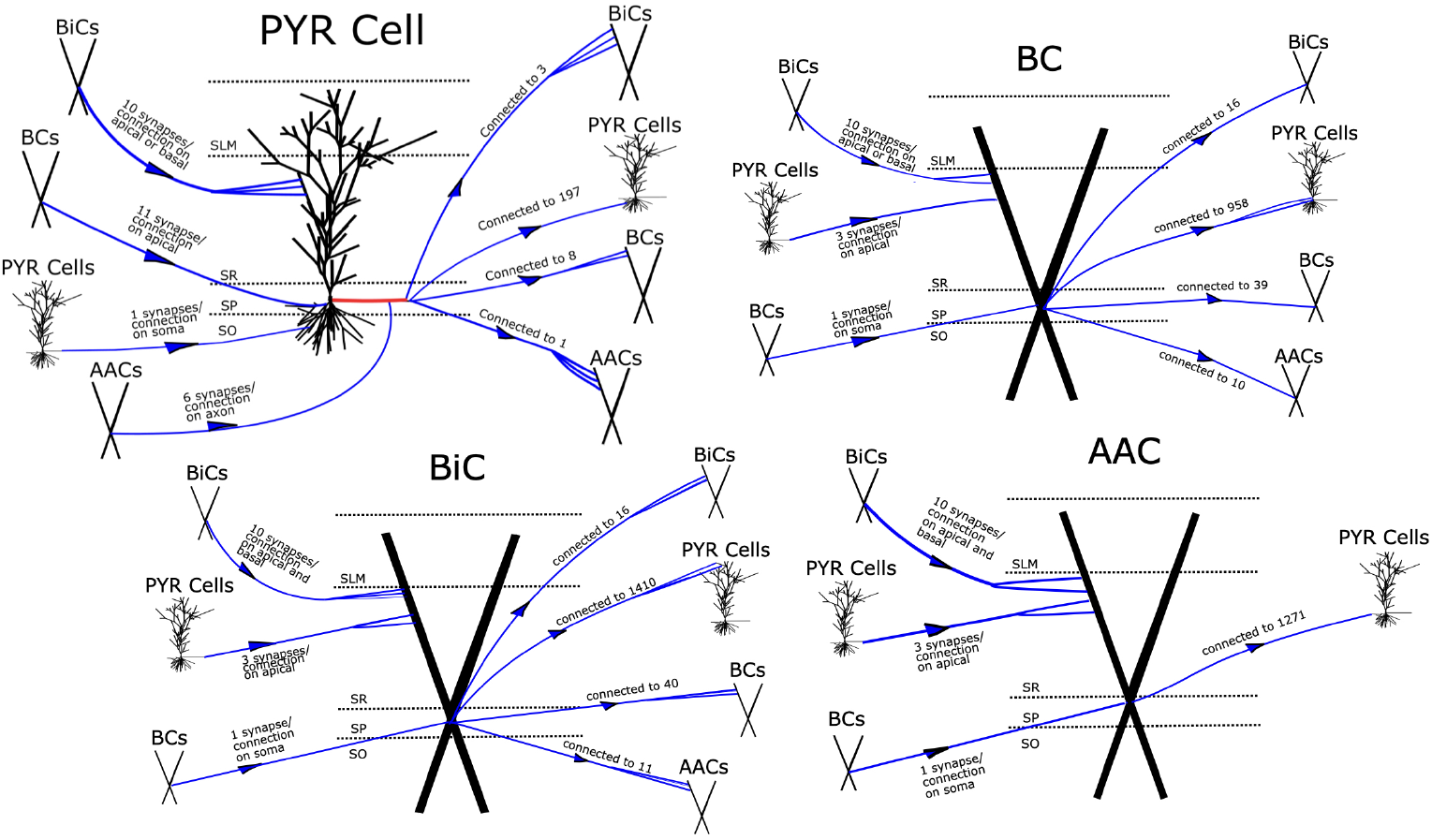
Schematics summarizing connections in the detailed model for PYR and PV+ cell types. The four schematics illustrate the connection schemes that exist in the detailed model, where we only consider PYR and PV+ cells (BCs, BiCs, AACs) (***Bezaire et al., 2016b***). For each large centered cell, the number of synapses per connection and its approximate location on the cell is specified for whichever cells are presynaptic, and the number of cells that the large centered cell connects to is also illustrated for whichever cells are postsynaptic. These numbers are also reflected in ***Table 4***. The morphological structure along with its layer location from the detailed model is also shown. The red line in pyramidal cell denotes its axon. SP = stratum pyramidale, SLM = stratum lacunosum-moleculare, SR = stratum radiatum, SO = stratum oriens. Other acronyms are defined in the main text.

The connection probabilities computed from the detailed model are given in ***Table 4*** along with connection probabilities from the minimal model (details are given in the Methods). To avoid repetition, minimal model connection probabilities are only shown for the “PV+=BC” case in row #2 of ***Table 4***. We found that regardless of the PV+ cell type consideration (i-iv), the connection probability from PV+ to PYR is greater than from PYR to PV+in the detailed model, indicating that one of the predictions of the minimal model is in effect in the CA1 microcircuitry. Thus, this comparison arguably yields a ‘validation’ of the minimal model as one of its main predictions is in effect in the detailed model which has empirically determined connection probabilities from many experimental determinations (***Bezaire and Soltesz, 2013***). We note that comparison of PYR to PYR and PV+ to PV+ connection probabilities between minimal and detailed models are expected to be appropriate as these connection probabilities in the minimal model were derived from the experimental literature (***Ferguson et al., 2013, 2015a***). As noted in ***Table 4***, the PYR to PYR connection probability (see row #1) is an order of magnitude less than it is for the PV+ to PV+ connection probability (see rows #2-#5) for both minimal and detailed models.

**Table 4.** Detailed Model Connection Probabilities and Synaptic Weights.

| Row | Cell Types and Connections | Number of cells | Number of connections | Connection Probability | Number of synapses per connection | Synaptic Weight*<br>*g(nS)* |
| --- | --- | --- | --- | --- | --- | --- |
| #1 | **PYR**<br>PYR to PYR | 311,500 | 197 | 0.00063<br>[0.01 for<br>minimal model (MM)] | 1 | 70.0<br>[0.094 for MM] |
| #2 | **PV+ = BC**<br>PYR to BC<br>BC to PYR<br>BC to BC | 5,330 | 8<br>958<br>39 | 0.0015<br>[0.02 for MM]<br>0.0031<br>[0.3 for MM]<br>0.0071<br>[0.12 for MM] | 3<br>11<br>1 | 2.1<br>[3.0 for MM]<br>2.2<br>[8.7 for MM]<br>1.6<br>[3.0 for MM] |
| #3 | **PV+ = BC/AAC**<br>PYR to BC/AAC<br>BC/AAC to PYR<br>BC/AAC to BC/AAC | 7,000 | 9<br>1,115<br>49 | 0.0014<br>0.0036<br>0.0070 | 6<br>8.5<br>1 | 4.4<br>5.7<br>0.8 |
| #4 | **PV+ = BC/BiC**<br>PYR to BC/BiC<br>BC/BiC to PYR<br>BC/BiC to BC/BiC | 7,740 | 11<br>1,184<br>111 | 0.0014<br>0.0038<br>0.014 | 6<br>10.5<br>11 | 16.0<br>3.7<br>77.1 |
| #5 | **PV+ = BC/AAC/BiC**<br>PYR to BC/AAC/BiC<br>BC/AAC/BiC to PYR<br>BC/AAC/BiC<br>to BC/AAC/BiC | 9,210 | 12<br>1,213<br>132 | 0.0013<br>0.0039<br>0.014 | 9<br>9<br>11 | 23.8<br>5.6<br>54.0 |
| #6 | **Other Input**<br>CA3 to PYR<br>EC to PYR<br>CA3 to BC<br>CA3 to AAC<br>EC to AAC<br>CA3 to BiC<br>EC to BiC | n/a<br>n/a<br>n/a<br>n/a<br>n/a<br>n/a<br>n/a | 5,985<br>1,299<br>6,047<br>4,170<br>485<br>5,782<br>432 | n/a<br>n/a<br>n/a<br>n/a<br>n/a<br>n/a<br>n/a | 2<br>2<br>2<br>2<br>2<br>2<br>2 | 0.40<br>0.40<br>0.44<br>0.24<br>0.24<br>0.30<br>0.30 |

\* Synaptic Weight = Synaptic Conductance × number of synapses/connection

In making these comparisons, we do not expect to have an exact matching of connection probability values. Besides the fact that the minimal model consists of a subset of different inhibitory cell types in the detailed model, the cellular models differ in their compartmental and mathematical biophysical ‘structure’. Specifically, the detailed model has multi-compartment models that include conductance-based ion current representations, and the minimal model has single compartment models with an Izhikevich mathematical representation (see Methods). It is however reassuring that the connection probabilities compare favourably as described above, since both minimal and detailed models produce intrinsic, intra-hippocampal theta rhythms.

#### E-I balance considerations in minimal and detailed models expose differences

So far we have shown that the connection probabilities in the minimal model are appropriate relative to the empirical ones in the detailed model and that the detailed model has appropriate EPSC/IPSC amplitude ratios from the perspective of the whole hippocampus preparation that generates intrinsic theta rhythms. Let us now exploit these linkages.

We first note that since both the minimal and full-scale detailed models produce theta rhythms, the underlying E-I balances that are present in both models must be appropriate for the generation of theta rhythms. Now, besides connection probabilities between excitatory and inhibitory cells, synaptic weights and any other external drives to the network models would also affect E-I balances.

##### Synaptic Weights

Similar to the comparison consideration of connection probabilities above, we compare synaptic weights in minimal and detailed models. As before, we focus on a cellular subset of the detailed model the fast-firing PV+ cells. The number of connections and synaptic weights for PV+ and PYR cells are given in the last two columns of ***Table 4***. Note that the synaptic weight refers to a connection between cells so that the number of synapses per connection is taken into consideration. From a comparison of these weights, it is clear that there is about three orders of magnitude difference between the synaptic weights of PYR to PYR cells whereas the synaptic weights from PV+ to PYR, PYR to PV+ and PV+ to PV+ are comparable (i.e., same order of magnitude), if PV+ cells are considered to be BCs or a combination of BCs and AACs (see ***Table 4***). Thus, on the face of it, the detailed model has much stronger connections between PYR cells relative to the minimal model.

##### External Drives

The minimal model is driven by an external excitatory input, denoted as ‘other input’ in the schematic of ***Figure 1***, that is applied only to the PYR cells of the E-I networks. The amount of this other input is comparable or smaller than any of the ‘internal’ EPSCs (see Table 5 in ***Ferguson et al. (2017)***), as it has a zero mean with fluctuations of ≈ 10-30 pA. For the detailed model, the excitatory and inhibitory cells are driven by activation of excitatory afferents from the CA3 and the entorhinal cortex (EC) with connectivity of empirical estimation (see row #6 in ***Table 4***). Unlike the minimal model, these CA3/EC excitatory inputs are larger relative to the ‘internal’ EPSCs and so likely play an important role in maintaining the appropriate E-I balance for theta generation in the detailed model. Specifically, the CA3, EC and PYR cell excitatory currents onto PYR cells are approximately 10, 6 and 10 nA. The detailed model is only loosely based on the whole hippocampus preparation. Its theta rhythms are produced intra-hippocampally but the network is driven by external EC and CA3 noisy afferents. These afferents conceptually represent remaining inputs from cut afferents after extraction from the whole brain. Given that the external drives in the minimal and detailed models are not represented in a similar way, we cannot compare them directly. However, it is possible that the large difference in PYR to PYR synaptic weights between minimal and detailed models is partly because of their external drive differences.

In summary, our consideration of linkages between minimal and detailed models via the whole hippocampus preparation (see ***Figure 1***) that generates intrinsic theta rhythms leads to the following: The minimal model has appropriate connection probabilities relative to the biological system, as represented by a biologically detailed full-scale CA1 microcircuit model; the full-scale detailed model has appropriate EPSC/IPSC amplitude ratios relative to experiment; and although both minimal and detailed models produce intra-hippocampal theta rhythms, there are notable differences between their PYR to PYR synaptic weights and external drives.

### Using a ‘piece’ of the detailed model to understand the initiation of theta rhythms and how their frequencies are controlled

It is worth re-stating that despite its several limitations (e.g., only 70% of inhibitory cell types were included), the detailed model produces robust theta rhythms. However, because of its large size and computationally expensive nature, extensive parameter explorations were not performed. As a result, even though the detailed model produces theta rhythms, and model perturbations indicated that some cell types and not others are important for their emergence, we do not know how the rhythm generation is initiated or controlled. To address this, we first isolate a part of the detailed model, the segment model (see ***Figure 1***), that has comparable cell numbers to the minimal model. We investigate the segment model according to the noted differences with the minimal model and examine how this is manifest in the power and frequency of LFP theta rhythms that we subsequently interpret in light of the minimal model mechanism. From this investigation, we unveil an understanding of how the ‘biophysical’ theta rhythms are generated and how their frequencies are controlled in a biologically detailed model with multiple inhibitory cell types.

#### Creating the segment model and examining its initial behaviour

We start by extracting a ‘piece’ of the detailed model which has a comparable number of cells relative to the minimal model, and we refer to it as the segment model - see ***Figure 8***Ai. Our segment model represents 10% of the original detailed model and it has all of the same cell types with the same layer location positioning and synaptic connection structure as the detailed model. That is, the segment model contains eight inhibitory cell types and is driven by excitatory afferents representing inputs from the EC and the CA3 region, as illustrated in ***Figure 8***Aii. The activation of the EC/CA3 synapses is modeled as an independent Poisson stochastic process and the strength of this activation is represented by the Poisson stimulation parameter. These afferents project to the majority of the cell types in the network with the exception of the OLM cells which are only driven by the PYR cells. Therefore, in contrast to the minimal model, the segment model is driven by external inputs that in addition to the PYR cells, also project to the inhibitory cells of the network (see ***Figure 1***). Even though the segment model represents only 10% of the original detailed hippocampus model, its much smaller size makes it now possible to investigate the network dynamics by undertaking extensive parameter explorations using high-performance computing. We carried out this investigation by exploiting the noted differences between minimal and detailed models, and by considering the minimal model insights.

**Figure 8.**
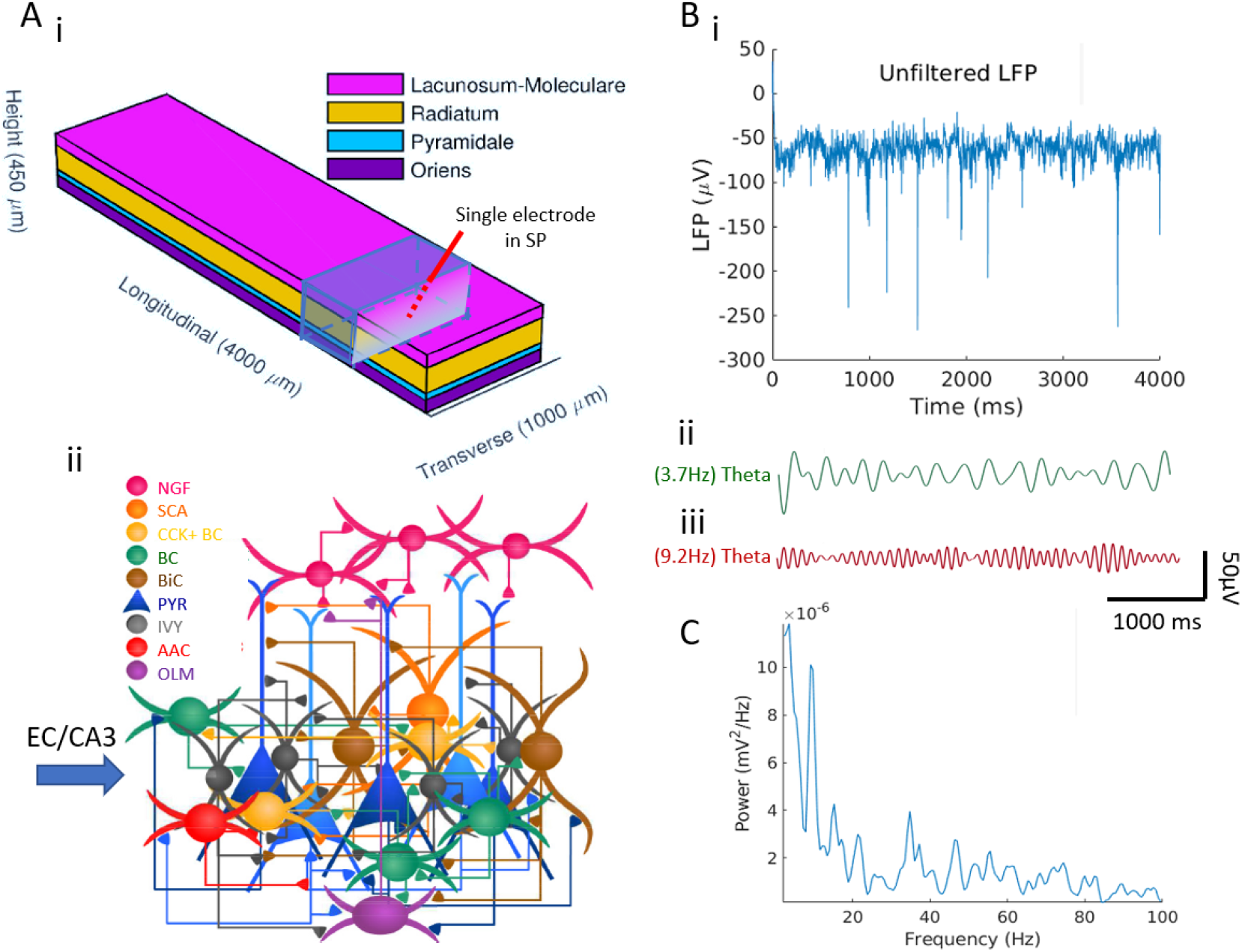
Theta rhythms in the segment model. **A**. (i): The model network is arranged in a layered prism. Image is adapted from Fig. 1 of ***Bezaire et al. (2016a)***. The segment model shown in blue, represents 10% of the original volume. It contains 31,150 PYR cells, 553 BCs, 221 BiCs, 358 NGF cells, 40 SCA cells, 360 CCK+ BCs, 881 Ivy cells, 164 OLM cells, 147 AACs. LFP output is based on a single micro-electrode placed in Stratum Pyramidale (SP). (ii): The number, position and cell types of each connection are biologically constrained, as are the numbers and positions of the cells. Image is adapted from Fig. 1 of ***Bezaire et al. (2016a)***. **B**. The segment network generates theta rhythms once the stimulation is reduced to 0.26Hz (it is 0.65Hz in the original detailed model). (i): Unfiltered LFP, (ii): filtered for low theta (peak at 3.7Hz) and (iii): filtered for high theta (peak at 9.2Hz). See Methods. **C**. Welch’s Periodogram of the LFP shows a peak at two theta frequencies. Acronyms are defined in the main text. **Figure 8-Figure supplement 1**. *Recurrent excitation and feed-forward external drive to the PYR cells are needed for theta rhythms*.

Let us start by examining the segment model without changing any of its parameters relative to the detailed model. As expected, the segment model does not produce any output. Instead, this ‘fraction’ of the detailed model produces hyperactive cell populations (not shown) indicating that the E-I input balances to the cells are shifted in favour of excitation. This suggests that to get a theta rhythm in the segment model, one could simply reduce the activation of the external afferents via the stimulation parameter. This is a reasonable consideration given that our model essentially consists of a smaller piece of tissue. We found that theta rhythms arise in the segment model when we decrease the stimulation parameter, but they have very low power and are very noisy. The raw LFP signal, as recorded in stratum pyramidale, is shown in ***Figure 8***Bi, and it can be seen to be quite noisy. Guided by the Welch’s Periodogram, as shown in ***Figure 8***C, theta rhythms at two peak frequencies (3.7 and 9.2 Hz) can be discerned. The filtered LFP signal is shown above ***Figure 8***Bii and Biii. In essence, this finding predicts that a 10% piece of a whole hippocampus preparation is enough of a tissue volume to generate theta rhythms. This supports the viewpoint, supported by experimental observations, that the hippocampus is comprised of multiple theta oscillators along its septotemporal axis (***Goutagny et al. (2009)***).

#### Designing an extensive parameter exploration of the segment model

As shown above, the segment model, without any changed parameter values besides the stimulation parameter, produces weak and noisy theta rhythms - see ***Figure 8***B. Is it possible to obtain robust theta rhythms in the segment model? That is, can we increase the power of the theta rhythms expressed by the segment model? To answer this, we were motivated to determine whether bringing the segment model to a similar E-I parametric regime as the minimal model could ‘enhance’ the theta rhythms. To test this, we examined whether by adjusting for differences between the models, we could increase the power of the theta rhythms expressed by the segment model.

From the comparison between the minimal and detailed models, we found that their two main differences stemmed from the external drives to the network and the synaptic weights between the PYR cells, which we will refer to as *g_pyr–pyr_*. In the minimal model, the external drive is only applied to the PYR cell population and is relatively weak (fluctuations of ≈ 10-30 pA) compared to what it is in the detailed model - about 10 nA (similar for the segment model). Also, the external drive in the detailed and segment models is applied not only to the PYR cells but also to the majority of the inhibitory cells. It is also important to keep in mind that the PYR cells in the segment model are bombarded by substantially more inhibition in comparison to the minimal model, as there are eight different inhibitory cell types projecting to them, as compared to just the fast-firing PV+ cells in the minimal model. This means that in the segment model, relative to the minimal model, it is possible that the stronger external drive to the PYR cells and the stronger *g_pyr–pyr_* are required to counterbalance the larger inhibitory presence due to the multiple inhibitory cell inputs. Due to these aspects, we designed an expansive exploration of how the segment model depends on *g_pyr–pyr_* and the external drive to the PYR cells in creating theta rhythms. For the external drive, we explored both the stimulation parameter as well as the excitatory conductance from EC/CA3 to the PYR cells, which we will refer to as *g*_*ec*/*ca*3–*pyr*_. This examination is schematized in ***Figure 9***A.

**Figure 9.**
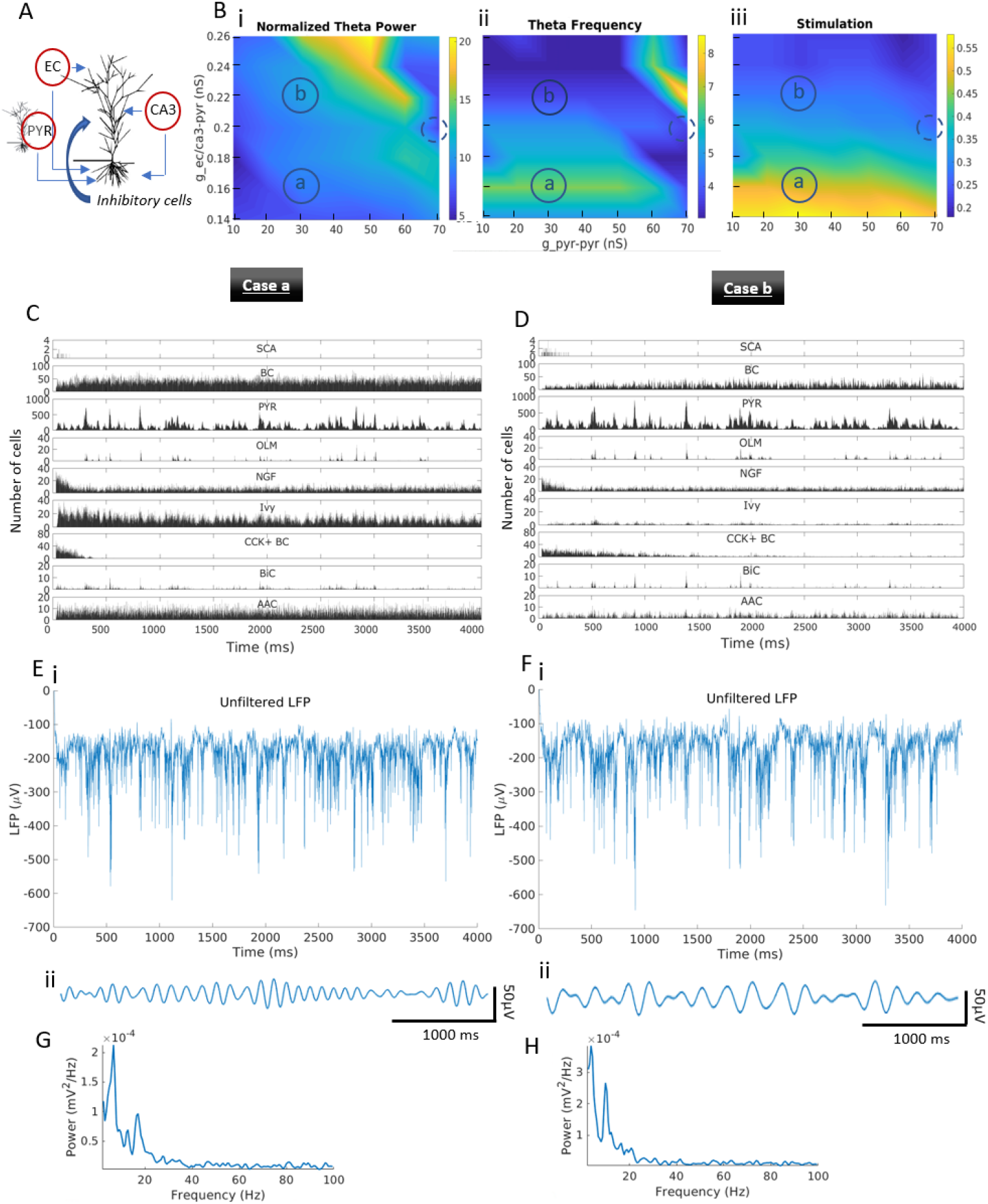
Dependence of theta power and frequency on the PYR cells’ excitatory drives. **A**. Schematic to illustrate the parametric exploration done that focuses on the excitatory drives to the PYR cells. **B**. Heatmaps of normalized theta power (i), frequency (ii) and afferent input stimulation (iii) as a function of *g_pyr-pyr_* and *g*_*ec*/*ca*3–*pyr*_. Circled a and b regions represent *case a* and *b* networks respectively, with (*g_pyr–pyr_*, *g*_*ec*/*ca*3–*pyr*_) parameter values of: (30 nS, 0.16 nS) for *case a*, and (30 nS, 0.22 nS) for *case b*. Dashed circled regions represent initial network of the segment model as obtained from the 10% ‘piece’ extracted from the detailed model (see ***Figure 8***), with (*g_pyr–pyr_*, *g*_*ec*/*ca*3–*pyr*_) parameter values of: (70 nS, 0.20 nS). **C**. Histograms of cellular activities for *case a*. Bin size = 1 ms. **D**. Same as C., but for *case b*. **E**. (i): Unfiltered LFP, (ii): Filtered LFP (peak at 6.7Hz), for *case a*. **F** (i): Unfiltered LFP, (ii): Filtered LFP (peak at 3.7Hz), for *case b*. **G**. Welch’s Periodogram of LFP for *case a*. **H**. Same as G., but for *case b*. **Figure 9-Figure supplement 1**. *Dependence of net theta power on the PYR cells’ excitatory drives*. **Figure 9-Figure supplement 2**. *Dependence of theta and delta power on the PYR cells’ excitatory drives*. **Figure 9-Figure supplement 3**. *Dependence of “high” theta (6-12Hz) power on the PYR cells’ excitatory drives*.

For each (*g_pyr–pyr_*, *g*_*ec*/*ca*3–*pyr*_) conductance pair, we performed a set of simulations to find the stimulation parameter that maximizes the theta power (3-12 Hz) for the given conductance pair. Given that these networks exhibit two theta peaks, a low and a high one, as shown by their Welch Periodogram, this analysis considers the stronger theta peak which is usually the one corresponding to the lower theta. A separate analysis for the higher theta peak power vs conductance pairs (ranges 6-12 Hz) can be found in ***Figure 9-Figure Supplement 3***. The theta rhythm dependence of our parametric explorations is shown in ***Figure 9***Bi-iii. From left to right we show the normalized theta power, the theta frequency and the required stimulation to maximize the theta power for each conductance pair examined. These results show that the normalized theta rhythm power increases with increasing *g_pyr–pyr_* or *g*_*ec*/*ca*3–*pyr*_ (similar is the trend for the net theta power ***Figure 9-Figure Supplement 1***) while theta frequency approximately decreases with increasing *g*_*ec*/*ca*3–*pyr*_ or *g_pyr–pyr_*. We note that these patterns are disrupted for the largest *g*_*ec*/*ca*3–*pyr*_ or *g_pyr–pyr_* conductance values, where the power of the networks is shifted to lower ‘delta’ frequencies below 3 Hz (see ***Figure 9-Figure Supplement 2***). From the heatmaps of the net theta power in ***Figure 9-Figure Supplement 1*** we notice that the power of the theta rhythms has significantly increased, approximately doubled, relative to the initial behaviour of the segment model shown in ***Figure 8***B. It is thus clear that there are particular parameter combinations that can significantly increase the power of the theta rhythms in the segment model to make it more robust.

#### Theta rhythm robustness and degeneracy of theta rhythm generation

To get an understanding of what underlies the results from our extensive parameter explorations, we took a detailed look at the inner mechanics of the network. We did this by examining two sets of conductance pair examples, *case a* (***Figure 9***C,E,G) and *case b* (***Figure 9***D,F,H), which correspond to small and large *g*_*ec*/*ca*3–*pyr*_ values, respectively. These two examples exhibit elevated theta power relative to the initial behavior of the segment, which we notice by comparing the amplitudes of the raw LFP recordings in ***Figure 9***Ei,Fi to ***Figure 8***Bi, and the periodograms in ***Figure 9***G,H to ***Figure 8***C, where the theta power can be seen to be larger by about two orders of magnitude. From our explorations, we observed the following: When *g*_*ec*/*ca*3–*pyr*_ is small, the EC/CA3 afferents have to be strongly activated to elicit a strong response to the PYR cells, hence requiring a large stimulation value - see ***Figure 9***Biii. However, because these afferents connect to most of the inhibitory cells, a large stimulation value means strong concurrent activation of most of the inhibitory cells in the network. This is why the majority of the inhibitory cells in the network are fairly active in these regimes as shown in ***Figure 9***C. When *g*_*ec*/*ca*3–*pyr*_ is large, the activation of EC/CA3 afferents don’t have to be as strong (see corresponding stimulation value in ***Figure 9***Biii) to elicit a similar response of the PYR cells given that the *g*_*ec*/*ca*3–*pyr*_ itself is already large. In this regime, the activity of most inhibitory cells is low exactly because the stimulation parameter is low and the inhibitory cells are not strongly activated. This can be seen in ***Figure 9***D.

Overall, these results expose the degeneracy of the theta rhythm-generating system which can occur in at least two ways depending on the exact pathway of activation of the PYR cells. It can be by either by low activation of the external afferents given a large *g*_*ec*/*ca*3–*pyr*_ conductance value, inducing a high concurrent activation of the inhibitory cells (*case a*), or by high activation of the external afferents given a small *g*_*ec*/*ca*3–*pyr*_ conductance value, inducing low concurrent activation of the inhibitory cells (*case b*). From this exploration, it is clear that regardless of the exact pathway of activation, what appears to be critical for robust theta rhythms is the net amount of input to the PYR cells. Thus, the proposition brought forth by the minimal model that the theta frequency is controlled by the net amount of input that is received by the PYR cells, seems likely. With the segment model, we are now in the position to directly examine whether this is the case.

#### Frequency control of theta rhythms and how they are initiated

Based on the minimal model’s proposition, we examined the frequency of the LFP theta rhythms from the perspective of the net current received by the PYR cells irrespective of whether the pathway is of a *case a* or of a *case b* type. To do this, we took advantage of the numerous network simulations underpinning the heatmaps of ***Figure 9***B. Specifically, we examined whether the frequency of those networks correlate with the net current to the PYR cells. We selected a sample of 10 PYR cells from each of the segment models, as schematized in ***Figure 10***A, and computed the average and standard deviation of the net current that each of these 10 PYR cells received. An example of IPSCs and EPSCs received by a particular PYR cell is shown in ***Figure 10***Bi-ii. In ***Figure 10***C, we plot means and standard deviations of the net current for all of the segment model networks in ***Figure 9***B, and we see that there is indeed a strong correlation between the theta frequency of each segment model and the net input received by the PYR cells (see Methods for calculations). This plot clearly demonstrates that the frequency of the theta rhythm can be predicted by the input to the PYR cells.

**Figure 10.**
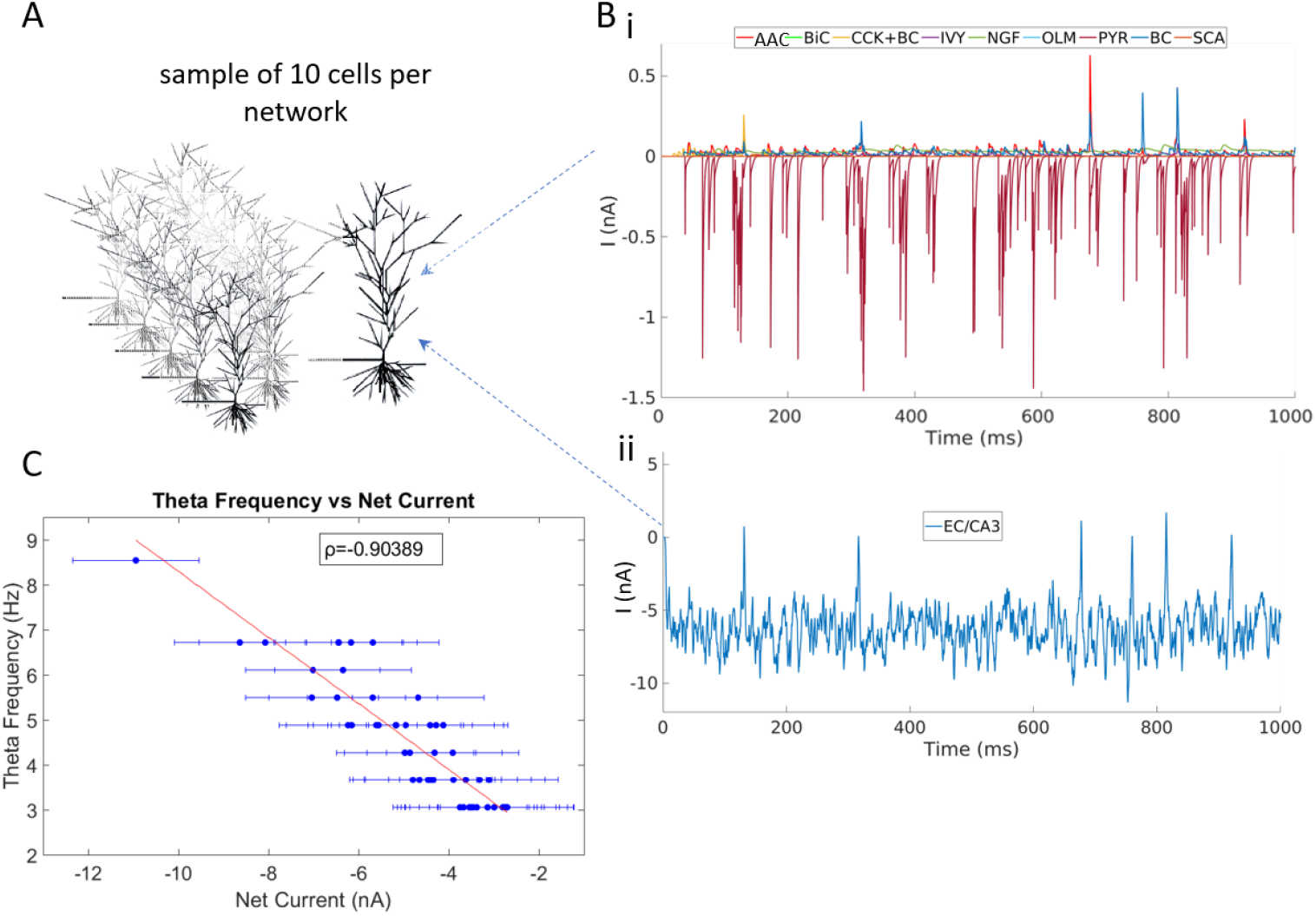
PYR cell net current input strongly correlates with frequency. **A**. Schematic to illustrate PYR cell sampling considered for net current analyses. **B**. Illustration of EPSCs and IPSCs onto the PYR cells. (i): current inputs from other PYR cells and the eight inhibitory cell types, and (ii): the excitatory drive from EC/CA3. **C**. Theta frequency plotted versus net current. Ten cells are randomly selected from each one of the 50 networks underpinning the heatmaps of ***Figure 9***B. Each dot represents the average across ten cells of the mean input current amplitudes to a given PYR cell of one of the 50 networks in ***Figure 9***B. Error bars represent the standard deviation of these averages. The correlation coefficient between the theta frequency and the net input current is *ρ* = −0.9, the p-value = 5.9×10^−^19 and the slope of the red line of the linear regression fit is r=-0.7 Hz/nA, indicating that the LFP theta frequency increases by about one Hz every time the net drive increases by one nA. Acronyms are defined in the main text.

So far we’ve shown that the frequency of the theta rhythm relies on the net input received by the PYR cells in the segment model representing the smallest volume of tissue required to produce theta rhythms. Indeed if we chose to consider an even smaller tissue volume some of the inhibitory cells wouldn’t even be part of the network purely because of their empirically derived connectivity profiles. At this point, we note that the presence of theta rhythms requires that PYR cells are connected with each other, since the rhythms do not exist if *g_pyr–pyr_* conductances are zeroed (see ***Figure 8-Figure Supplement 1***). That is, some recurrent excitation is required, as was already shown in ***Bezaire et al. (2016b)***. Also, not surprisingly, given the large contribution of the external drive in the detailed model, the theta rhythm cannot be maintained if external drive to the PYR cells is removed by setting *g*_*ec*/*ca*3–*pyr*_ to zero (see ***Figure 8-Figure Supplement 1***). Interestingly, what becomes evident in the segment model is that the generation of the theta rhythms is not specifically due to phasic drives from the inhibitory cells. Indeed, in these networks most of the inhibitory cell populations haven’t yet organized into periodically firing populations. This is particularly noticeable in ***Figure 9***D where theta rhythms are present and can be seen to be due to the PYR cell population firing in bursts of theta frequency. Even more, we notice that the pattern of the input current to the PYR cells isn’t theta-paced or periodic (see ***Figure 10***Bi). Despite this, the PYR cell population can organize into a theta frequency bursting population, and initiate the theta rhythm. This indicates that provided the appropriate level of net input to the PYR cells, a theta rhythm can start, and the initiation does not depend upon sequential, externally imposed inhibition form other rhythmically firing inhibitory cells. Of course, with a larger network, other inhibitory cells organize into periodically firing populations and contribute to the robustness and strength of the theta rhythm. However, at its initiation stages, we can clearly say that the theta rhythm ‘sparks off’ from the PYR cells.

#### Experimental constraints expand the understanding of theta-generating mechanisms in the hippocampus

Given the not unexpected degeneracy in the segment model, an important aspect to consider is which of the theta rhythm-generating pathways might be occurring in the biological system. As a step in this direction, we turn to experimental observations from the intact hippocampus in which PV+ cells were optogenetically manipulated by ***Amilhon et al. (2015)***. Specifically, it was found that optogenetically silencing the PV+ cells significantly reduced the theta rhythm. Thus, removing PV+ cells in the segment model should have a detrimental effect on theta rhythms as well. As already noted, there are several sets of parameters that produce theta rhythms, and these are shown in ***Figure 9***B.

Let us go back to our previous examples of *case a* and *case b*. As can be seen in ***Figure 9***G,H, these two networks produce theta rhythms of similar power. To consider the experimental results of ***Amilhon et al. (2015)***, we removed the PV+ cells (BCs, AACs, BiCs) from the two network cases to mimic an ‘optogenetic’ silencing, and we measured the resulting change in the theta rhythm. This was done by removing the PV+ cells from the network by zero-ing all of the inhibitory synaptic conductances emanating from them (***Figure 11***A, ***Figure 11***B-G). It is evident that the PV+ cell removal has a negative effect on the power of the theta rhythms in *case a* but not in *case b*, simply based on their respective periodograms (compare ***Figure 11***F,G with ***Figure 9***G,H). Interestingly, there was a large increase in gamma frequencies with PV+ cell removal in *case a*. In *case a*, the net input to the PYR cells is the sum of both strong inhibitory and excitatory currents; thus, the rhythm cannot be maintained when the inhibitory inputs from PV+ cells are lost due to the severe disruption of the E-I balance. However, in *case b*, the net input to the PYR cells is mostly defined by the excitatory cells. In this case, removing the PV+ cells did not affect the E-I balance enough to disrupt the theta rhythms - indeed, it enhanced them (compare the peak values in the periodograms of ***Figure 9***H and ***Figure 11***G). This implies that the different E-I balances in the segment model that allow LFP theta rhythms to emerge are not all consistent with the experimental data, and by extension, the biological system. Thus it appears that lower *g*_*ec*/*ca*3–*pyr*_ conductance values, as in *case a*, that rely on both inhibitory and excitatory currents are more consistent with the experimental data.

**Figure 11.**
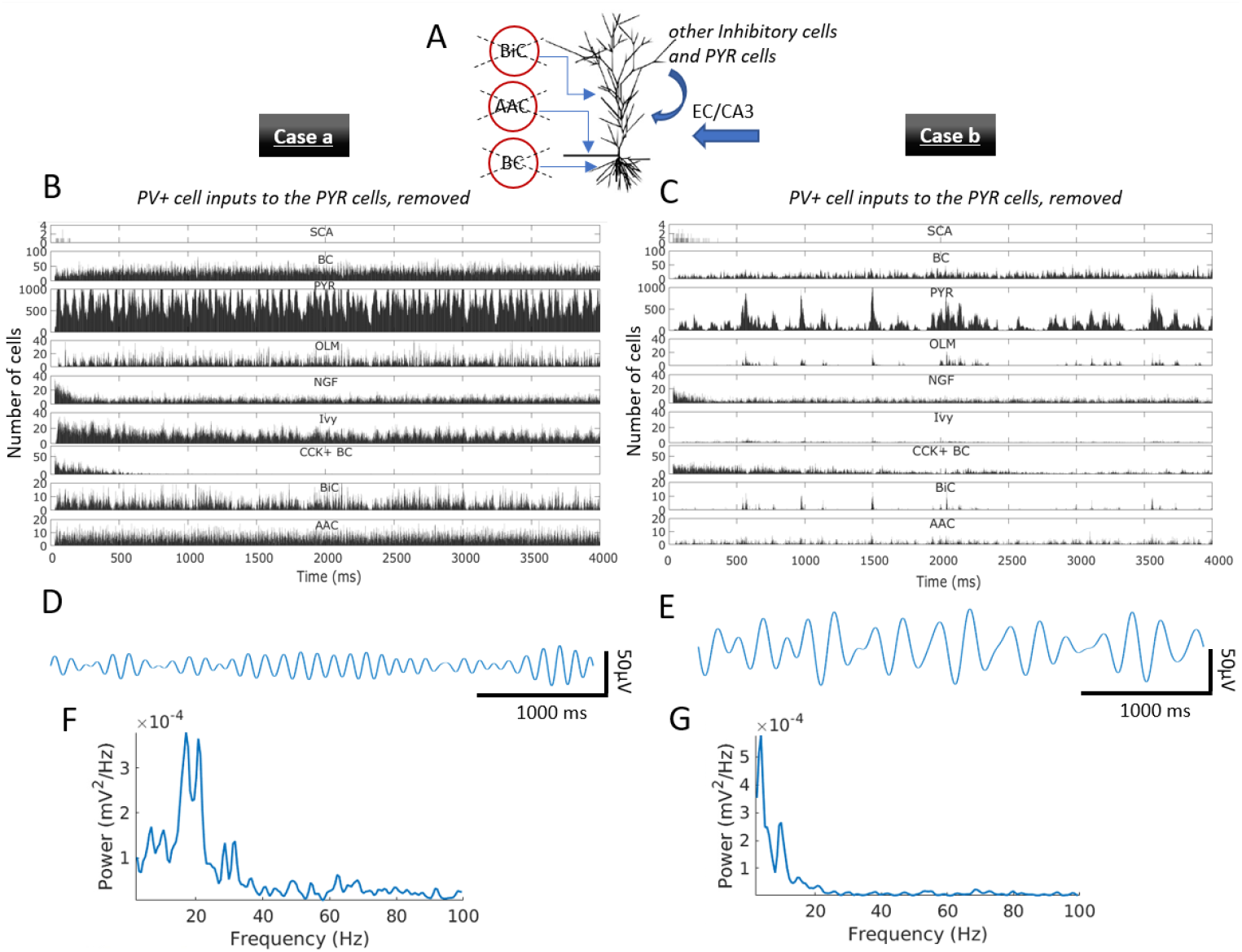
Effect on the theta rhythms with removal of input from PV+ cells. **A**. Schematic illustrating examination of the effects of PV+ cell (BCs, AACs, BiCs) input removal to the PYR cells. **B**. Histograms of cellular activities for *case a* with PV+ to PYR cell inputs removed. Bin size = 1ms. **C**. Same as B., but for *case b*. **D**. Filtered theta signal for *case a* with PV+ to PYR cell inputs removed (peak at 6.7Hz). **E**. Same as D., but for *case b* (peak at 3.7Hz). **F**. Welch’s Periodogram of LFP for *case a* with PV+ to PYR cell inputs removed. **G**. Same as F., but for *case b*. Acronyms are defined in the main text.

In ***Figure 12*** we show a summarized, aggregate comparison of the measurements for *case a* and *case b* segment models before and after the removal of the PV+ cells from the network. In *case a* (***Figure 12***Ai-iv), removing the PV+ cells diminishes the theta power, while the frequency of the LFP signal and the net input current to the PYR cells which are correlated, remained intact. A noticeable decrease appears in the standard deviation of the current. This decrease reveals that removing the PV+ cells in this regime increases the ‘noisiness’ of the net current, or the fluctuation around its mean, which could potentially underlie the decrease in theta power in this example. Indeed, after examining the minimal model in the first part of this study, we proposed an ‘inhibition-based tuning’ mechanism for the theta rhythm, in which the PV+ cells ‘tune’ the PYR cell firing and by consequence regularize and enhance the robustness of the theta rhythm. Such a mechanism is supported by the segment model for *case a*.

**Figure 12.**
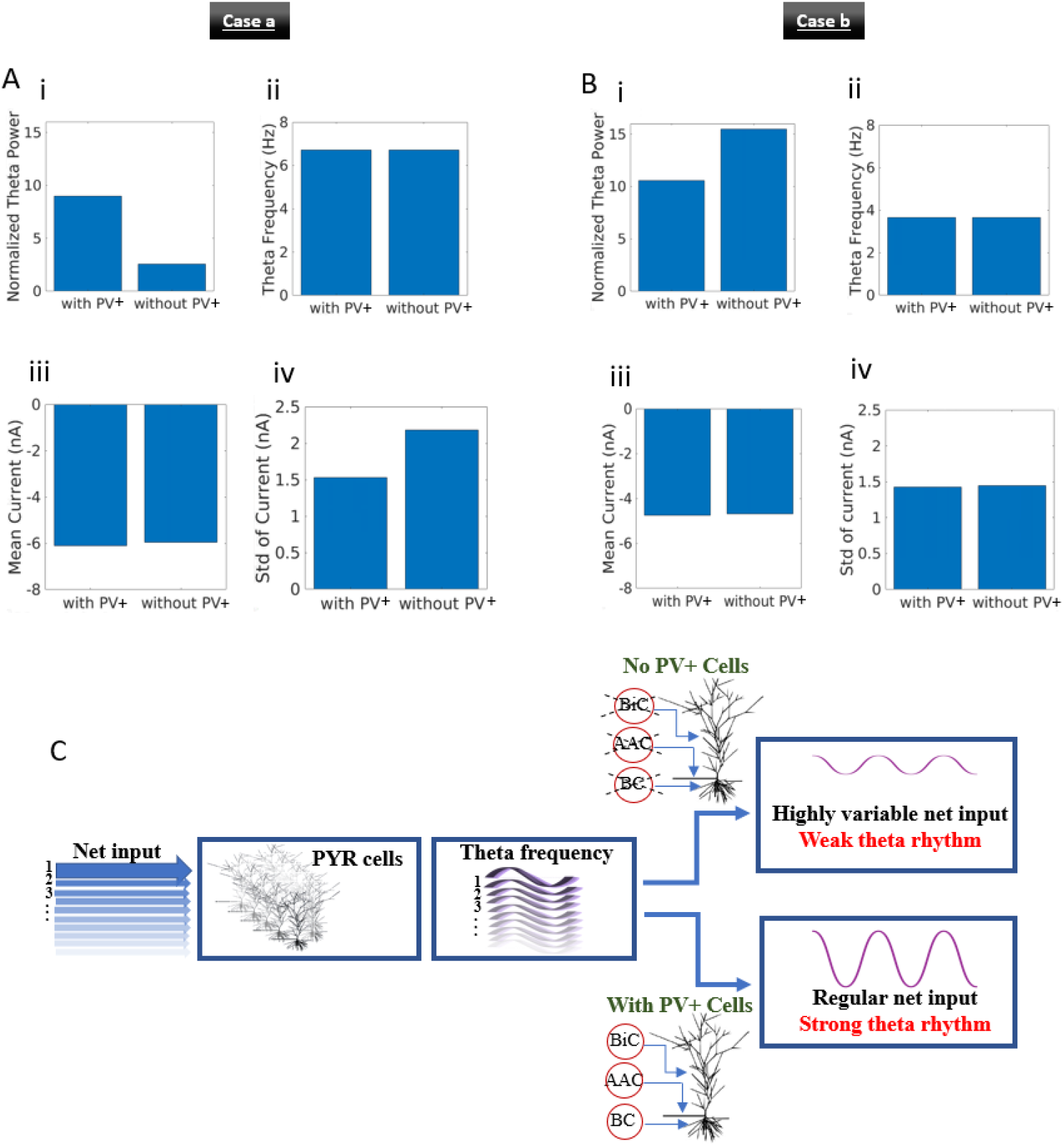
Aggregate comparison of theta rhythms before and after the removal of inputs to PYR cells from PV+ cells and schematic of ‘biophysical theta’. **A**. Results for *case a*. (i): Normalized theta power, (ii): theta frequency, (iii): mean current, and (iv): standard deviation of current, with and without PV+ cells for *case a*. **B**. Same as **A**., but for *case b*. **C**. The net PYR cell input controls the resulting theta frequency. The PV+ cells contribute to the net input while they also regularize it and amplify theta power.

As shown for *case b* (***Figure 12***Bi-iv), removing the PV+ cells actually increases the power of the theta rhythm while keeping the same theta frequency in the LFP signal and the same net input current. However, in this case, the standard deviation of the net current did *not* change, unlike for *case a*. Thus, from the perspective of the experiments of ***Amilhon et al. (2015***) theta rhythm generation via a *case a* type pathway seems more biologically realistic while it also supports the proposed inhibition-based tuning mechanism from the minimal model. In ***Figure 12***C, we provide a schematic of the biophysical theta generation mechanism and frequency control. This comparison with experiment brings forth the importance of understanding the inner mechanisms underpinning the dynamic output of a system, as high-dimensional models are likely to express degeneracy, which could however come forth via separable “pathways” of different biological implications.

## Discussion

Including biological complexity in cellular-based network models challenges our ability to understand their dynamic behaviours. To tackle this challenge, we have brought together two previously published models of the CA1 microcircuit that generate theta rhythms without oscillatory inputs. The two models mimic the intrinsic theta rhythms of an intact, whole hippocampus preparation (***Goutagny et al., 2009***). One of them - the minimal model (***Ferguson et al., 2017***) - only has fast-firing PV+ and PYR cells, whereas the other - the detailed model (***Bezaire et al., 2016b***) - has eight different inhibitory cell types and PYR cells. The minimal model uses a simplified Izhikevich mathematical model structure for cellular representations, with parameter values determined from fits to experimental data from the whole hippocampus preparation, whereas the detailed model uses multi-compartment conductance-based cellular representations, determined from an extensive knowledge-based review of the literature (***Bezaire and Soltesz, 2013***).

The wide variety of cell types that make up brain circuits leads to high-dimensional sets of nonlinear, differential equations described by large sets of parameters incorporated into models. This makes application of theoretical analyses difficult and parametric explorations computationally expensive. In our approach of bringing together the two models in this study, we implemented a focused, hypothesis-driven parametric search of a fragment of the detailed model, the segment model, guided by the minimal model. This allowed us to establish a cellular basis for how intrinsic theta rhythms are generated and how their frequencies are controlled in CA1 microcircuits of the hippocampus. The importance of considering multi-level and multi-granular networks to understand brain phenomena as done here, was recently discussed by ***Einevoll et al. (2019)***.

### Summary overview

We started from the minimal model where it was previously shown that population bursts of theta frequency can be generated in E-I networks with sparse firing of PYR cells and EPSC/IPSC current amplitude ratios as observed experimentally. This occurred due to *SFA, Rheo* and *PIR* building block features. Using heterogeneous PYR cell populations and quantification of *SFA, Rheo* and *PIR* building block features, we explored the robustness of the theta generation mechanism in the minimal model and found that it is sensitive to specific *Rheo* and *PIR* quantified values, but not to *SFA*. We subsequently used PRCs to determine how the frequency of theta rhythms could be controlled, and proposed an ‘inhibition-based tuning’ mechanism in which inhibitory inputs to the PYR cell population allow a stable theta rhythm to emerge, given an appropriate net input to the PYR cells. This paved the way for investigations with the detailed model where this could be directly examined.

Since the detailed model was not explicitly built with the whole hippocampus preparation in mind, we computed EPSC/IPSC amplitude ratios and confirmed that they were in line with those observed experimentally in the whole hippocampus. Comparisons between minimal and detailed models validated the predicted connectivity balance in the minimal model and exposed notable differences.

We extracted a ‘piece’ of the detailed model of comparable cell numbers as the minimal model - termed the segment model - and showed that it could generate theta rhythms, albeit noisy and of low LFP power. This finding supports the experimental observations of ***Goutagny et al. (2009***) that the theta rhythm in the whole hippocampus is composed of a set of coupled oscillators, and only a part of the entire hippocampus is required to generate theta rhythmic output, an ‘oscillator’. With this smaller segment model, we focused our investigation on the differences between the minimal and the detailed model, namely the PYR-PYR synaptic weights and the external drives.

We found a strong correlation between the theta oscillation frequency and the average net input delivered to the PYR cells. This indicates that the frequency of the LFP theta rhythm can be predicted by the inputs to the individual PYR cells of the network. Further investigations of the segment model revealed that the theta rhythm is initiated by the PYR cells but is regularized by the PV+ cells since their removal caused a large decrease in the LFP power and an increase in the variability of the net current received by the PYR cells. Together, this supports an inhibition-based tuning mechanism for theta generation (see ***Figure 12***C).

### Mechanism underpinnings and leveraging of theoretical insights

From our previous work we already knew that minimally connected PYR cell networks produced theta frequency population bursts on their own (***Ferguson et al., 2015a***), but the majority of the PYR cells would fire during population theta bursts which is unlike the experimental observations of sparse PYR cell firing. With the inclusion of PV+ cells to create E-I networks, the population of PYR cells fired sparsely, which makes sense since the addition of inhibitory cells leads to less firing of PYR cells due to silencing from the inhibition. Relatedly, it has been shown that feedforward inhibition plays a role in maintaining low levels of correlated variability of spiking activity (***Middleton et al., 2012***).

It is important to point out different PYR cell aspects in the minimal and detailed models. As mentioned, for the minimal model we know that the PYR cell population on its own can generate a population theta rhythm, and this is by virtue of its intrinsic properties that includes an *SFA* building block feature (***Ferguson et al., 2015a***). In that previous work, we had used a PYR cell model that is strongly adapting based on fits to the experimental data, or weakly adapting based on another experimental dataset in the same paper (***Ferguson et al., 2015b***), that could produce theta frequency population bursts in both cases. As discussed in ***Ferguson et al. (2015b)***, it is unlikely that there are distinct types of biological PYR cells that are strongly or weakly adapting, but rather a continuum of adaptation amount dependent on the underlying balances of biophysical ion channel currents. Our explorations of the robustness of the theta generation mechanism in the minimal model revealed that theta rhythms are not sensitive to the specific quantified value of the SFA building block feature, so long as there is some adaptation. Thus, although the minimal model from ***Ferguson et al. (2017***) used a strongly adapting PYR cell model and the mimimal model database used here started from this strongly adapting PYR cell model basis, it is unlikely that our results would be affected.

For the detailed model, the PYR cell model is based on experimental data in which some adaptation can be seen in the experimental recording, but is not apparent in the PYR cell model output of the detailed model (see Appendix of ***Bezaire et al. (2016b)***). This then suggests that the prediction of the segment model that the PYR cells are the initiator of theta rhythms is not simply due to adaptation. It must thus involve other intrinsic characteristics of the the biophysical PYR cell models. That excitatory networks can produce population bursts in of themselves is not new to the theoretical, modeling world, but it has not been previously shown that this could be the case in a biophysically detailed CA1 microcircuit model. An important candidate among PYR cell intrinsic properties that affect *PIR* is the hyperpolarization-activated (h-) channel (***Ascoli et al., 2010***). The h-channel has been shown to be a pacemaking current and contributes to subthreshold resonance (***Biel et al., 2009***). It has been a focus in general network modeling studies (e.g., ***Avella Gonzalez et al. (2015)***), as well as specific to inhibitory cells in the generation of coherent oscillations (***Rotstein et al., 2005***). It is interesting to note that the h-channel, with its non-uniform distribution, has been shown to play an important role in shaping the output of LFP recordings, as determined from multi-compartment LFP modeling studies (***Ness et al., 2016***, ***2018***; ***Sinha and Narayanan, 2015***). How exactly h-channels in PYR cells influence the dynamics and frequency of LFP theta rhythms in CA1 microcircuits will be interesting to investigate further.

As shown in our heterogeneous PYR cell E-I network explorations, the presence of theta rhythms (i.e., population bursts in the minimal model) was sensitive to the specific quantified values of *PIR* and *Rheo* building block features. It is expected that there would be a sensitivity to *Rheo* as the rheobase current of PYR cells dictate whether a PYR cell would spike or not. We had noted that an Izhikevich cellular model requires a positive *b* value in order for *PIR* to occur - i.e., for a spike to fire after hyperpolarization, and while there is sensitivity to this *PIR* value, it is not the case that PYR cell firing occurs on rebound from inhibition during the ongoing theta population bursts (see ***Figure 4***). In actual CA1 PYR cells, it has been shown that *PIR* spiking does occur, mediated by h-channels, and is locally controlled by biophysical ion channel balances (***Ascoli et al., 2010***). Whether PYR cells actually fire due to *PIR* during ongoing theta rhythms may or may not be the case, and one could potentially disentangle this in the model with consideration of the variety of inhibitory cell types. However, this seems less critical to figure out now that we have exposed a strong correlation between the frequency of the theta rhythm and the net current to the individual PYR cells. We know that *PIR* is present in CA1 PYR cells, and we know that the minimal model indicates it as a sensitive feature for theta rhythms, and we thus predict that changes to the PYR cell’s intrinsic properties that affect *PIR* would affect the resulting theta rhythms.

PRC theory has been used in a variety of ways in the neuroscience field (***Schultheiss et al., 2011***), and particularly in consideration of network dynamics. For example, ***Hansel et al. (1995***) used PRCs to explain the differential capacity for excitatory signalling to synchronize networks of Type I or Type II neurons (these types are differentiated by their bifurcation type (***Izhikevich, 2006***)), ***Rich et al. (2016)*** analyzed synchronization features in purely inhibitory networks using PRCs, and ***Achuthan and Canavier (2009)*** used PRCs to understand clustering in networks. We took advantage of PRC theory by considering phase-resetting of the PYR cells in the E-I networks due to incoming inhibitory input. In this way, we were able to hypothesize an inhibition-based tuning mechanism for control of the theta rhythm frequency based on the PRC shape (amount of advance or delay) and the PYR cell’s intrinsic firing frequency. Our use of PRCs relied on our observations of the effect of different PRC shapes on the resulting theta rhythm. Such a consideration is similar to that used by ***Rich et al. (2016)*** to explain differential synchrony patterns in inhibitory networks of Type 1 vs Type II neurons.

### Physiological considerations and related studies

Based on the number of cells, the minimal and segment models are designed to represent a ‘piece’ of CA1 microcircuitry, and not the whole hippocampus preparation. However, the ability of these models to generate population theta rhythms on their own, is in line with the observations of ***Goutagny et al. (2009)*** where transmission between portions of the whole hippocampus preparation were blocked with procaine (see their supplementary Fig.11). With each piece of tissue being able to generate theta oscillations on its own, the whole hippocampus would represent a set of coupled oscillators. Indeed, traveling theta waves in hippocampus and neocortex have been considered in this fashion (***Lubenov and Siapas, 2009***; ***Zhang et al., 2018***). In previous work, we used phase-coupled oscillator models, assumed inhibitory coupling between oscillators and examined asymmetries in coupling strengths that could be responsible for the experimentally observed propagation of slow rhythms (***Skinner et al., 2001***). In that vein, it may be worth considering whether one could combine the mechanistic insights from microcircuit and coupled oscillator model studies.

The extensive set of simulations performed with the segment model showed that different cell-specific pathways dominate LFP theta rhythms of similar frequency and power, exposing degeneracy. While model degeneracy in high-dimensional model systems is expected, it underlines the importance of probing generation mechanisms whenever possible, and not just comparing outputs. There are multiple pathways in the circuitry, and at the *in vivo* level, one cannot unambiguously disentangle these pathways or have cell-type considerations (***Benito et al., 2014***). Using the segment model, we were able to consider two distinct ‘pathways’ by which theta rhythms are generated - one where the EC/CA3 to PYR cell inputs dominated (*case b*) and another where they did not (*case a*). Based on perturbative responses to the model to mimic the experiments, only *case a* was in accordance with experimental data (***Amilhon et al., 2015***). We note that the differences between the cases could actually reflect differences in the contributions of particular inhibitory populations since, for example, the recordings that we compare our simulations to are taken from the superficial layers of the hippocampus. Indeed, in a very recent modeling study by ***Navas-Olive et al. (2020)*** that built on the detailed model of ***Bezaire et al. (2016b)***, it was shown that deep and superficial PYR cells fire at different phases of the theta oscillation and are driven by different inhibitory cell populations. In that study, the authors found that in CA1, PV+ BCs preferentially innervate PYR cells at the deep sublayers while CCK+ BCs are more likely to target superficial PYR cells. It is possible thus, that our *case b* regime reflects a theta rhythm relevant to the deep CA1 layers which is highly modulated by the CCK+ BCs, which, in contrast to the PV+ BCs, happen to be particularly active in *case b*. However, what is clear from our work is that specific perturbations could determine the dominance of different cellular pathways by comparing LFP output characteristics.

The determination of an inhibition-based tuning mechanism for theta generation stemmed from this study is essential, as it forms a foundation from which to consider E-I ‘balances’ during theta rhythms from detailed physiological and experimental perspectives. E-I balances have been shown to be quite precise in feedforward networks from CA3 to CA1 (***Bhatia et al., 2019***), and fine-scale mapping studies show structured synaptic connectivity between different cell types in these regions (***Kwon et al., 2018***). Thus, in the absence of a detailed enough cellular-based network model one could not really situate emerging biological details’ contributions to theta rhythms. On the other hand, in the absence of some mechanistic understanding, the importance of various biological details is challenging to contain. In this work, we have combined the strengths of minimal and detailed models, and have perhaps reached an ‘inflection point’ (***Gjorgjieva et al., 2016***) by having enough, but not too much, biological realism to obtain a cellular-based mechanistic understanding. Had we started from models that were either more abstract or more detailed, model linkages and mechanism translations may have not been possible (i.e., too far from an ‘inflection point’).

### Limitations and future work

Even though our modeling study sheds light on the foundation of the theta mechanism, more can still be unveiled in terms of the specific roles of the variety of inhibitory cell types in the segment model and their inter-relationships. Through optogenetic perturbations, experimental studies have already explored how PV+ as well as somatostatin-positive (putative OLM cells) cells affect intra-hippocampal theta rhythms (***Amilhon et al., 2015***). Our previous modeling work examined the contribution of BiCs, BCs and OLM cells to ongoing theta rhythms and LFP generation (***Chatzikalymniou and Skinner, 2018**; **Ferguson et al., 2015c***) in light of these experimental studies. However, the segment model, with its complement of eight inhibitory cell types and its computational tractability, provides an exciting opportunity to extract and predict specific inhibitory pathways and their activation machinery during theta rhythms. Achieving this will help guide and target perturbation and stimulation paradigms in pathological states.

Besides ***Bezaire et al. (2016b)***, other detailed CA1 microcircuit models that include multiple inhibitory cell types have been developed ((***Cutsuridis et al., 2010***; ***Shuman et al., 2020***; ***Turi et al., 2019***)). However, these models were used to examine higher level behaviours and theta rhythms were imposed, not generated within the models. Recently, a very detailed quantification of synaptic anatomy and physiology that includes short-term plasticity has been done, and is provided as a resource for the community (***Ecker et al., 2020***). It may be possible to examine these other detailed models in light of our mechanistic understanding, and further, to design a strategy that would appropriately include additional inhibitory cell types in the CA1 microcircuit model via the determined mechanism.

### Concluding remarks and a proposal: A ‘pacemaker circuit’

Six years ago, Siegle and Wilson’s work (***Siegle and Wilson, 2014***) showed strong support for phase coding in the hippocampus, using the encoding and retrieval paradigm developed by Hasselmo (***Hasselmo et al., 2002***) with theta rhythms. Recognizing the multi-layered aspects of theta rhythms - different cholinergic sensitivities, distinct phase relationships with different inhibitory cell types, low and high frequency theta types, different behavioural correlates and information processing, dorsal and ventral differences, heavy dependence on medial septal circuitry interactions (***Chauvière, 2020***; ***Colgin, 2013, 2016***; ***Hinman et al., 2018***) - our work plants a seed.

Until now, it was not clear how one could consider theta rhythms from both cell-type pathways with E-I balances and functional behavioural perspectives. Our work suggests that there is no longer a need to separately impose theta rhythms on network models, as the cells in these networks are themselves part of the theta rhythm-generating machinery and this ‘separation’ eliminates some of the interactions that may be critical and thus hinder our understanding of the system. What is clear is that there *is* a theta rhythm generator in the hippocampus, i.e., intrinsic theta rhythms can be generated in a whole hippocampus preparation (***Goutagny et al., 2009***). We know that interactions with the medial septum (MS) are important for theta, but we note that lesioning the MS reduces, but does not terminate theta rhythms (***Colgin, 2013***; ***Winson, 1978***). Modeling work has suggested that theta rhythms could arise due to hippocampo-septal interactions (***Hajós et al., 2004***; ***Wang, 2002***). It is likely that interactions with the MS circuitry act to make the intrinsic hippocampus theta rhythms more robust, and impose theta rhythms in MS. Interestingly, experimental data has shown that rhythmic stimulation of the hippocampo-septal fibers can ‘phase’ MS neurons at that exact frequency due to rebound dependent h-channels, suggesting that the intrinsic hippocampus theta generator could be transferred to MS neurons via E-I interactions (***Manseau et al., 2008***). At present, we are not aware of any evidence supporting that the MS can generate theta rhythms on its own.

Thus we propose that CA1 PYR cells act as theta rhythm initiators tuned by the inhibitory cell populations to create a ‘pacemaker circuit’ - a core theta generator - in the hippocampus, with PYR cells sensitively dependent on ‘pacemaking’ h-channels. Amplification of these rhythms occurs due to inputs from the MS, while the net input received by the PYR cells controls the resulting theta frequency. From this intrinsic theta rhythm foundation, we can build, and in the process, disentangle the cellular-based and multi-layered aspects of theta rhythm generation and function in the hippocampus (***Brandon et al., 2011***; ***Koenig et al., 2011***; ***Jaramillo and Kempter, 2017***), and possibly other brain structures, since interestingly, functional connectivity studies have shown that the hippocampus is a brain hub (***Battaglia et al., 2011***; ***Mišić et al., 2014***). A schematic of our proposal is shown in ***Figure 13***.

**Figure 13.**
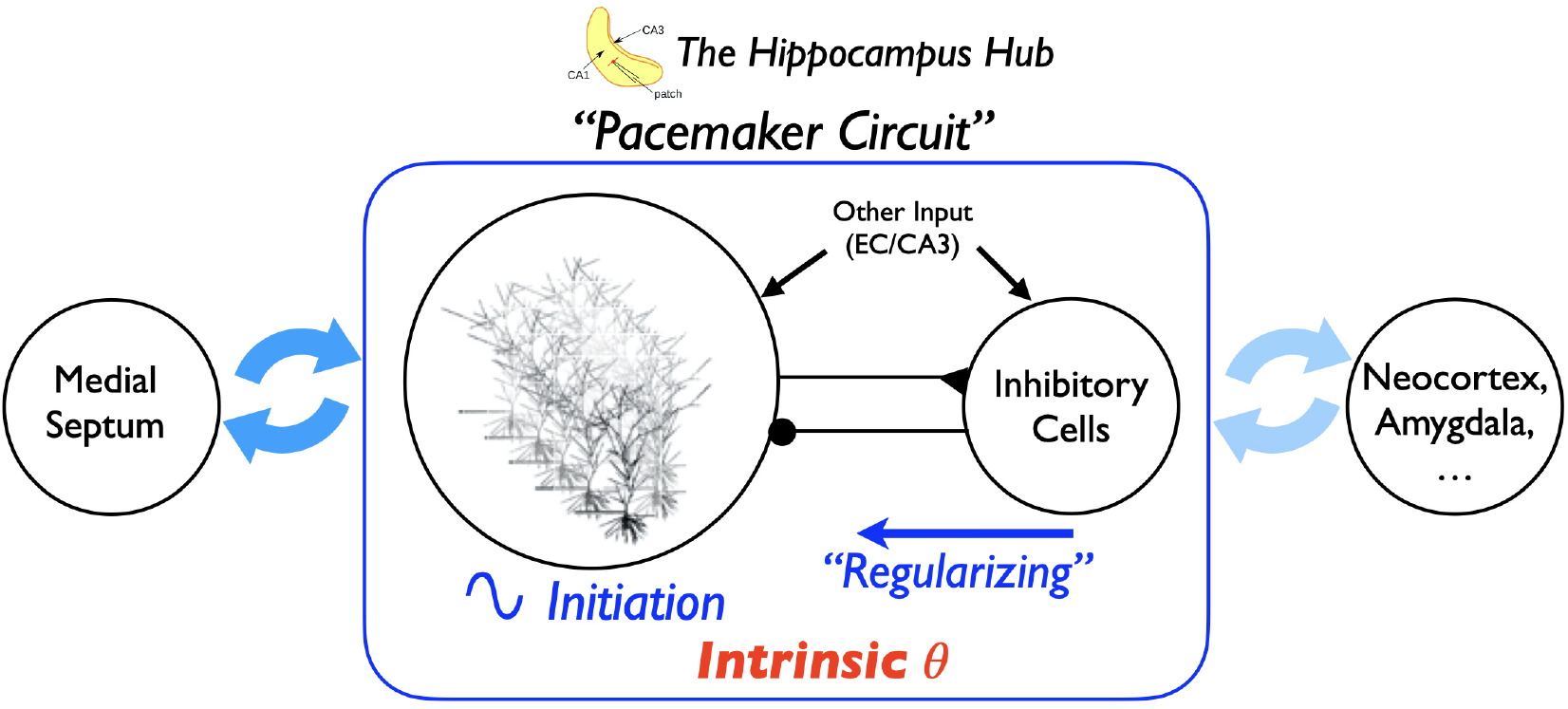
Proposing a theta pacemaker circuit in a hippocampus hub. The hippocampus can produce intrinsic theta oscillations on its own, without the need for any oscillatory input. In the work here, we have shown that theta rhythms can be generated by the PYR cell population, and are ‘tuned’ and regularized by the inhibitory cell population, as illustrated in the rectangle. We propose that this theta pacemaker circuit is amplified by connections with the MS via hippocampo-septal cellular interactions, as illustrated by the dark blue thick arrows. That is, the MS is not a theta rhythm generator, but rather acts to enhance and amplify the existing intrinsic theta rhythm in the hippocampus, and would play a role in setting the particular theta rhythm frequency. This would occur due to the MS cellular inputs affecting the net input current to the PYR cells in the hippocampus. The theta rhythm would further interact with other regions such as neocortex and amygdala, as illustrated by the light blue thick arrows (***Battaglia et al., 2011***). The possibility of a hippocampus hub is supported by connectivity studies (***Mišić et al., 2014***). The whole hippocampus schematic is adapted from Fig 1 of ***Huh et al. (2016)***.

## Methods

### The minimal model and expanded explorations

Details of the minimal model rationale and setup are previously published in ***Ferguson et al. (2017)***, but some background relevant to the present work is summarized here. The minimal model represents an approximate one mm^3^ ‘piece’ of the CA1 region of the hippocampus determined to be enough to generate theta rhythms (***Goutagny et al., 2009***). It has 30,500 cells (30,000 excitatory, PYR cells and 500 inhibitory, fast-firing PV+ cells). In analyses of excitatory networks on their own, a scaling relationship between cell number, connection probability and excitatory synaptic weight allowed us to use 10,000 PYR cells rather then 30,000 in the excitatory network simulations (***Ferguson et al., 2015a***). As the model is minimal, we could perform thousands of simulations on high-performance computing to ascertain parameter balances that would produce theta rhythms as well as capture experimental data results of EPSC/IPSC amplitude ratios. For this to be the case, we found that the connection probability from PV+ to PYR cells should be larger than from PYR to PV+ cells (***Ferguson et al., 2017***).

We note that the PV+ cells have intrinsic and synaptic connectivity aspects derived from experiment and that inhibitory PV+ cell networks fire coherently given appropriate excitatory drives and synaptic weights (***Ferguson et al., 2013***). In the E-I networks of the minimal model, the excitatory drive to PV+ cells comes from the PYR cell population (see schematic in ***Figure 1***). We note that when we did the E-I network simulations in ***Ferguson et al. (2017)***, we chose the synaptic weight (between PV+ cells) to be such that it could be at the ‘edge’ of firing coherently (high frequency) or not (see Fig. 3 in ***Ferguson et al. (2013)***). As such, given an appropriate excitatory drive, it can be switched into a high frequency coherent regime so that the PV+ cell network could produce an inhibitory ‘bolus’. From estimates of EPSCs onto the PV+ cells of 1000 pA, the synaptic weight between PV+ cells was set to 3 nS (***Ferguson et al., 2017***).

#### Cellular specifics and equations

The network structure and cellular details for the minimal model simulations in the present paper are similar to those in ***Ferguson et al. (2017)***. That is, cellular models (PYR and PV+ cells) are based on experimental data from the *in vitro* whole hippocampus preparation (***Ferguson et al., 2013, 2015b***). They use the mathematical model structure developed by Izhikevich (***Izhikevich, 2010***, ***2006***), in which the subthreshold behaviour and the upstroke of the action potential are captured, and a reset mechanism to represent the spike’s fast downstroke is used. Despite being relatively simple, parameter choices can be made such that they have a well-defined (albeit limited) relationship to the electrophysiological recordings. It has a fast variable representing the membrane potential, *V* (*mV*), and a variable for the slow “recovery” current, *u* (*pA*). We used a slight modification to be able to reproduce the spike width. It is described by the following set of equations:

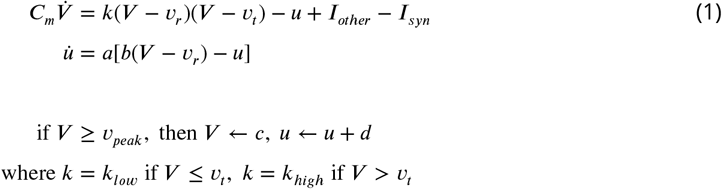

where *C_m_* (*pF*) is the membrane capacitance, *υ_r_* (*mV*) is the resting membrane potential, *υ_t_* (*mV*) is the instantaneous threshold potential, *υ_peak_* (*mV*) is the spike cut-off value, *a* (*ms*^−1^) is the recovery time constant of the adaptation current, *b* (*nS*) describes the sensitivity of the adaptation current to subthreshold fluctuations - greater values couple *V* and *u* more strongly resulting in possible subthreshold oscillations and low-threshold spiking dynamics, *c* (*mV*) is the voltage reset value, *d* (*pA*) is the total amount of outward minus inward currents activated during the spike and affecting the after-spike behaviour, and *k* (*nS/mV*) represents a scaling factor. *I_syn_* = 0 for the isolated cell. *I_other_* is as described below for computing metrics for the PYR cell or E-cell.

Model parameter values for the PV+ cell or I-cell (units above) are: *υ_r_*=-60.6; *υ_t_*=-43.1; *υ_peak_*=-2.5; *c*=-67; *k_high_*=14; *C_m_*=90; *a*=0.1; *b*=-0.1; *d*=0.1; *k_low_=1.7*. These parameters are as previously determined (***Ferguson et al., 2013***), and are not varied. Model parameter values (units above) for the PYR cell are: *υ_r_*=-61.8; *υ_t_*=-57; *υ_peak_*=22.6; *c*=-65.8; *k_high_*=3.3; *C_m_*=115; *a*=0.0012; *b*=3; *d*=10; *k_low_*=0.1. These parameters are as previously determined for strongly adapting cells (***Ferguson et al., 2015b***), and the *a, b, d, k_low_* parameters *are* varied.

#### Network specifics and equations

The cellular models described above were used to create excitatory-inhibitory (E-I) networks as done in ***Ferguson et al. (2017)***. Specifically, synaptic input between PYR cells (E-cells), PV+ cells (I-cells) and between PYR and PV+ cells by representing synaptic input in Equation 1 as:

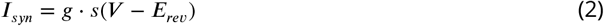

where *g*(*nS*) is the maximal synaptic conductance of the synapse from a presynaptic neuron to the postsynaptic neuron, *E_rev_*(*mV*) is the reversal potential of the synapse, and *V* (*mV*) is the membrane potential of the postsynaptic cell. The gating variable, *s*, represents the fraction of open synaptic channels, and is given by first order kinetics (***Destexhe et al. (1994)***, and see p.159 in ***Ermentrout and Terman (2010)***):

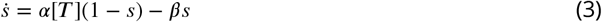

The parameters *α* (in *mM*^−1^*ms*^−1^) and *β* (in *ms*^−1^) in Equation 3 are related to the inverse of the rise and decay time constants (*τ_R_, τ_D_* in *ms*). [*T*] represents the concentration of transmitter released by a presynaptic spike. Suppose that the time of a spike is *t* = *t*_0_ and [*T*] is given by a square pulse of height 1 *mM* lasting for 1 *ms* (until *t*_1_). Then, we can represent

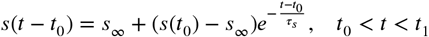

where 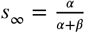 and 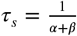. After the pulse of transmitter has gone, *s*(*t*) decays as

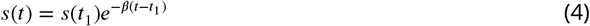

For network simulations, *I_other_*. in Equation 1 represents ‘other input’ to the PYR cell population (see ***Figure 1***), and is given by *I_other_* = −*g_e_*(*t*)(*V* – *E_rev_*). *g_e_*(*t*) is a stochastic process similar to the Ornstein-Uhlenbeck process as used by Destexhe and colleagues (***Destexhe et al., 2001***)

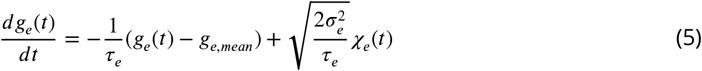

where *χ_e_*(*t*) is an independent Gaussian white noise process of unit standard deviation and zero mean, *g_e,mean_* (*nS*) is the average conductance, *σ_e_* (*nS*) is the noise standard deviation value, and *τ_e_* is the time constant for excitatory synapses. *τ_e_* is fixed based on values as used in ***Destexhe et al. (2001)*** (*τ_e_* = 2.73 *ms*).

Parameter values (rationale and refs given in ***Ferguson et al. (2017)***) are: *E_rev_*= −15 or −85 mV for excitatory or inhibitory reversal potentials respectively. Rise and decay time constants are, respectively, 0.27 and 1.7 msec for PV+ to PV+ cells; 0.3 and 3.5 msec for PV+ to PYR cells; 0.37 and 2.1 msec for PYR to PV+ cells; 0.5 and 3 msec for PYR to PYR cells. Connection probabilities are fixed at 0.12 for PV+ to PV+ cells and 0.01 for PYR to PYR cells, as estimated from the literature. For the simulations in this paper, we use connection probabilities that were found to be in line with the experimental data. That is, where the connection probability from PV+ to PYR cells (*c_PV,PY R_*) be larger than from PYR to PV+ cells (*c_PY R,PV_*).

Specifically, for the heterogeneous networks examined in this paper, we mainly focus on parameter values from Table 5 of ***Ferguson et al. (2017***)): *g_pyr_*=0.094 nS, *σ_e_*=0.6 nS, *g_pyr–pυ_*=3 nS, *g_pυ–pyr_*=8.7 nS, *c_PY R,PV_*=0.02, *c_PV,PY R_*=0.3, *g_e,mean_* = 0 nS. An actual instantiation of the ‘other input’ that these parameter values produce can be seen in the schematic figure of ***Figure 5***. We also consider networks with parameter values of: *g_pyr_*=0.014 nS, *σ_e_* =0.6 nS, *c_PYR,PV_*=0.02, *c_PV,PYR_*=0.3; and *g_pyr_*=0.084 nS, *σ_e_*=0.2 nS, *c_PYR,PV_*=0.04, *c_PV,PYR_*=0.5; and *g_pyr_*=0.084 nS, *σ_e_*=0.6 nS, *c_PYR,PV_*=0.02, *c_PV,PYR_*=0.5 (*g_pyr–pυ_*, *g_pυ–pyr_*, *g_e,mean_* the same as focused parameter values), and similar results are obtained. From the minimal model we know that theta population bursts occur when PYR cells receive zero mean excitatory drive with fluctuations of ≈ 10-30 pA (as estimated from 0.2 to 0.6 nS ‘noise’) (***Ferguson et al., 2017***).

#### PYR cell (E-cell) model database and building block feature quantifications

To create a database of PYR cell models, we range the *a, b, d, k_low_* model parameter values to create 10,000 models, 10 different values for each parameter, so as to encompass the default values from ***Ferguson et al. (2015b)*** obtained in creating the strongly-adapting PYR cell model based on experimental data from the whole hippocampus preparation. The default values of the strongly adapting PYR cell model are: *a*=0.0012*ms*^−1^; *b*=3.0*nS*, *d*=10p*A*, *k_low_*=0.10*nS/mV* and for the PYR cell model database, the parameter ranges are: [*initial value, final value, resolution*]: *a* = [0.0, 0.00216, 0.00024]; *b* = [0.0, 5.4, 0.6]; *d* = [0,18, 2]; *k_low_* = [0.0, 0.18, 0.02].

For each PYR cell model, spike frequency adaptation (*SFA*), post-inhibitory rebound (*PIR*) and rheobase (*Rheo*) building block features are quantified to allow comparisons to be made. The Euler integration method is used to integrate the cell equations with a timestep of 0.1 msec. Quantification of building block features is done as follows:

##### Rheo

Starting from *υ_r_*, each PYR cell model is given a constant current from −25 to 25 pA in 0.5 pA increments. If a spike is generated within the first 500 msec, then that constant current value is considered as the rheobase current, and is taken as the *Rheo* quantified value.

##### PIR

Starting from *υ_r_*, each PYR cell model is subjected to a one second hyperpolarizing step current for current values from 0 to −25 pA with a resolution of 0.5 pA. If a spike occurred upon termination of a given hyperpolarization step (i.e., a PIR spike) but not at the previous step value, then that step value is considered as the *PIR* quantified value.

##### SFA

Starting from *υ_r_*, each PYR cell model is subjected to input currents for one second, from 0 to 98 pA (inclusive) in 2 pA increments. For each input current, the number of spikes is recorded, and the interspike interval is calculated between the first and second spikes, and the last and second from last spike. The inverse is taken and defined as the initial and final frequency at that current. The initial and final frequencies as a function of the current steps creates a smooth, approximately linear relationship, so lines are fitted to the initial and final frequency plots. The slopes of those lines are subtracted from one another (the initial slope is always steeper) to produce the *SFA* quantified value.

The range of quantified values obtained from the model database of 10,000 PYR cells is: *SFA*: −0.001 to 0.64 (Hz/pA); *Rheo:* 1.5 to 6.5 (pA); *PIR*: −23.5 to −1.0 (pA). How they end up being distributed is shown in ***Figure 2***, and while clearly not a uniform or normal distribution, they encompass a wide range of values. The quantified values for the strongly adapting PYR cell model that we use as our starting basis in generating the model database (see above for full model and parameter values) are: *SFA*= 0.46; *Rheo*= 4.0; *PIR* = −5.0. We refer to them as the base values.

#### Heterogeneous PYR cell setup

The two ways in which heterogeneous PYR cell populations are created is as follows:

i. Using narrow (*N*) or broad (*B*) ranges of values for [*SFA, Rheo, PIR]* relative to base values, where *N* or *B* means that [*SFA, Rheo, PIR]* metric values are ± [0.1,0.5, 0.5] or ± [0.45, 3.0, 5.0] respectively, of base values. Thus, *NNN* refers to models with [*SFA, Rheo, PIR]* values of: [(0.36 to 0.56 exclusive of bounds; 4.0; −5.0], and *BBB* refers to models with [*SFA, Rheo, PIR]* values of: [(0.01 to 0.64 exclusive of bounds (noting that 0.64 is the maximum possible in the model database set); 1.5, 2.0, 2.5, 3.0, 3.5, 4.0, 4.5, 5.0, 5.5, 6.0, 6.5; −0.5, −1.0, −1.5, −2.0, −2.5, −3.0, −3.5, −4.0, −4.5, −5.0, −5.5, −6.0, −6.5, −7.0, −7.5, −8.0, −8.5, −9.0, −9.5]. Note that since the resolution of the *Rheo* and *PIR* quantified values are 0.5, and the manner in which it is defined (see above), the *N* range for *Rheo* has models in which *Rheo* = 4.0 only, and similarly, the *N* range for *PIR* has models in which *PIR* = −5.0 only. The other sets (using ranges as defined above) have quantified values as follows: *BBN*=[(0.01 to 0.64 exclusive of bounds; 1.5, 2.0, 2.5, 3.0, 3.5, 4.0, 4.5, 5.0, 5.5, 6.0, 6.5; −5.0]; *BNB*=[(0.01 to 0.64 exclusive of bounds; 4.0; −0.5, −1.0, −1.5, −2.0, −2.5, −3.0, −3.5, −4.0, −4.5, −5.0, −5.5, −6.0, −6.5, −7.0, −7.5, −8.0, −8.5, −9.0, −9.5]; *NBN*=[(0.36 to 0.56 exclusive of bounds; 1.5, 2.0, 2.5, 3.0, 3.5, 4.0, 4.5, 5.0, 5.5, 6.0, 6.5; −5.0]; and so on for *BNN, NBB*, and *NNB*. These eight possible cases and the number of models in each of them is given in ***Table 2***, along with the population frequency and power. Parameter value histograms for each of these combinations from the model database set are given in https://osf.io/yrkfv/, and what ranges of the quantified values in the database that they encompass is shown in ***Figure 2***.
ii. Using low (*L*), medium (*M*) or high (*H*) values, with *SFA* quantified value ranges exclusive of endpoints given as: *SFA*: *L* = [(0.0 to 0.2)], *M* = [(0.2 to 0.4)], *H* = [(0.4 to 0.6)]; *Rheo: L* = [1.5, 2.0, 2.5], *M* = [3.5, 4.0, 4.5], *H* = [5.5, 6.0, 6.5]. *PIR*: *L* = [−3.5, −4.0, −4.5], *M* = [−6.5, −7.0, −7.5], *H* = [−9.5, −10.0, −10.5]. This means that the base values fall into the *HML* case, with the small caveat that the *PIR* base value is just outside the *L* range. The gaps in these ranges are due to the automation of the exploration and to ensure that there is no overlap in the quantified values for a given case. Note that there ended up being no models for the cases: *HHH, HHL, MHH, MHL, LHH, LHL*, from the created model database. Thus there are 21 cases from the generated model database, and the number of models present in each case is given in ***Table 2***, along with population frequency and power. Parameter value histograms for eight of these cases are given in https://osf.io/yrkfv/, and what ranges of the quantified values in the database that they encompass is shown in ***Figure 2***.

#### E-I networks and simulations

To build E-I model networks, we choose PYR cells from the model database in two ways in consideration of *SFA*, *Rheo* and *PIR* building block features, referring to them as a trio in the following order: [*SFA, Rheo, PIR]*. The chosen PYR cells are distributed among the 10,000 cells to be used in the E-I network simulations in the following way: An individual PYR cell model is randomly chosen from the set of models of a particular heterogeneous PYR cell population that have [*SFA, Rheo, PIR]* values within the specif ed range. For example, if there are 33 PYR cell models in the set, then the number of cells conforming to each of the 33 PYR cell models should approach 10,000/33 in the E-I network, but there may not be an exactly equal number of the different PYR cell models. That is, we do the following: If there are 33 PYR cell models in the given heterogeneous PYR cell model set, then each PYR cell model out of 10,000 in the E-I network is given a random number between 1 and 33, and assigned that model’s parameters. We note that comparisons between the heterogeneous E-I networks are not perfectly ideal since the number of different PYR cell models varies (see ***Table 2***), and so the ‘amount’ of heterogeneity would vary in the various E-I networks. However, since we are mainly considering whether the theta rhythm would be lost or not, this is deemed to be acceptable.

The minimal model E-I network simulations are done using the Neuroscience Gateway (NSG) for high-performance computing (***Sivagnanam et al., 2013***). Simulations are run for 10 seconds using the Euler integration method with a timestep of 0.04 msec. The frequency and network power of the network simulation is computed as before (***Ferguson et al., 2017***). That is, for each network simulation, the population activity is defined as the average membrane potential of all the cells, with the frequency and network power taken as frequency and spectral peak from a fast Fourier transform (FFT) calculation of the population activity.

Code details are provided in https://github.com/FKSkinnerLab/CA1_Minimal_Model_Hetero and simulation output in https://osf.io/yrkfv/.

#### Phase response curve computation specifics

Phase response curves (PRCs) are calculated for each of the PYR cell models as described below. In Figure 6 the PRCs in each “model set” are averaged and presented along with a range of ± one standard deviation (shown by the shading around the curve).

Each PRC is calculated in the following fashion: A set input current (either 20 or 30 pA) is tonically applied to the cell, and the period (defined *λ*) of the cell’s firing is calculated as the time between the ninth and tenth cell spike. The inverse of the period represents the firing frequency of the cell, reported as averages and standard deviations for entire model sets in Figure 6. We calculate the phase response of the neuron to a perturbation at 100 equidistant times in its normal firing cycle. Here, the perturbation is a 1 ms current pulse with −500 pA amplitude. For 1 ≤ *i* ≤ 100, we define 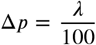 and deliver the perturbation at *i* * Δ*p* ms after the 10th cell spike. We then measure the time between the 10th and 11th cell spike as the “perturbed period” (defined *λ_p_*). We calculate the difference between this and the previously calculated period (in the absence of any perturbation) and normalize this by the normal firing period, meaning that in the PRC plots the y-axis is 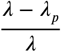. This means that negative values plotted in the PRC correspond with a phase-delay, i.e. the perturbed period was longer than the unperturbed period, and vice-versa. The x-axis in the PRC plots are the normalized time at which the perturbation was delivered, simply calculated as 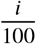. We note that we perform this calculation separately for each *i*, i.e. we re-initialize the cell and let it respond naturally to a tonic input until the 10th spike for each value of *i*, rather than perform these perturbations sequentially and risk confounding the responses.

The code for generating and plotting these PRCs can be found at https://github.com/sbrich/Theta_PRCs.

### The segment and detailed models and explorations

The segment model is simply a 10% piece of the detailed model of the rodent CA1 microcircuit (***Bezaire et al., 2016b***) as illustrated in ***Figure 1*** and ***Figure 8***A. To create and use the segment model, one must first be able to access and use the detailed model.

In segment and detailed models, there are eight different inhibitory cell types and excitatory PYR cells. All of these cell types are connected in empirically specific ways based on an extensive knowledge-based review of the literature (***Bezaire and Soltesz, 2013***). The cells are evenly distributed within the various layers of the CA1 (stratum lacunosum-moleculare, radiatum, pyramidale, oriens) in a three-dimensional prism. Afferent inputs from CA3 and EC are also included in the form of Poisson-distributed spiking units from artificial CA3 and EC cells. We note that although there are layer-dependent specifics regarding how the different cell types are arranged in the full-scale detailed model (***Figure 1***), there are not any differences along the longitudinal axis of the full-scale model. As such, the connection profile at any location along the longitudinal axis does not vary. In other words, the connection probabilities in any particular part of the longitudinal axis would be the same assuming that there are enough cell numbers for meaningfulness in the calculations.

#### Accessing the CA1 microcircuit model

The code that we use for this work starts from the original CA1 microcircuit repository which can be found at ModelDB at: https://senselab.med.yale.edu/ModelDB/showModel.cshtml?model=187604. The model version we used can be downloaded from: https://bitbucket.org/mbezaire/ca1/pull-requests/3/d1efeb957848/commits. Analysis of simulation outputs can be recreated using the publicly available SimTracker tool (***Bezaire et al., 2016a***) which can be downloaded from: http://mariannebezaire.com/simtracker/. It is recommended that users install SimTracker first and then install and register the ca1 model under SimTracker, to take advantage of the visualization functionalities of the SimTracker package. This tool is offered both as a stand-alone, compiled version for those without access to MATLAB (for Windows, Mac OS X, and Linux operating systems), and as a collection of MATLAB scripts for those with MATLAB access. Once the SimTracker and the ca1 repository are installed, users can run simulations either on their local machines using a small scale of the CA1 network, or on supercomputers as needed for full scale network simulations. To reproduce the findings presented here, one needs to first familiarize oneself with the CA1 microcircuit background and code.

The segment model is created from the detailed model by setting the “Scale” parameter = 10, which reduces the number of cells in the network by a tenth, and then dividing all connections in the network by a factor of 10. If this latter step is not done, then each cell would have ten times as many connections relative to a cell in the full-scale detailed network. That is, the parameter scaling is a ‘normalization’ in which the ‘scaled’ network assumes that each cell is a representative of ‘10 cells’. We did not want this, as the segment model is simply a piece of the detailed model and so we ‘removed’ the normalization by dividing the number of connections by ten.

#### Calculation of connection probabilities and synaptic weights in the detailed model

To be able to compare connectivities between minimal and detailed models, we compute connection probabilities in the detailed model. They are computed by dividing the total number of connections from a single presynaptic cell of a given type, to the cells of the postsynaptic population, divided by the total number of (postsynaptic) cells, of that particular population They are thus computed as divergent connection probabilities, as it was done in the minimal model where random divergent connection probablities were employed. To compute connection probabilities when PV+ cells are assumed to consist of more than one inhibitory cell type, a combination is required. For example, in considering BCs and BiCs as fast-firing PV+ cells in one population, the number of connections each cell (either BC or BiC) receives is the average of presynaptic connections each receives, as given in the detailed model. For example, the number of connections from PYR cells onto BC/BiC population equals the total number of presynaptic connections that BCs and BiCs receive from PYR cells. The connection probability from PYR to PV+ cells (BC/BiC combination) is calculated by dividing this total number of connections by the total number of BCs and BiCs. All numbers and connection probabilities are shown in ***Table 4***.

The synaptic weight in the detailed model is given by the synaptic conductance multiplied by the number of synapses per connection. So, for example, as a single BC cell has 11 synapses/connection onto a PYR cell and a synaptic conductance of 0.2 nS, then the synaptic weight is 2.2 nS. In the case of combined cell type populations, the average synaptic weight for the given cell type with its number of synapses/connection and synaptic conductance as reported by ***Bezaire et al. (2016b)***. All of the computed synaptic weights are shown in ***Table 4***.

#### Calculation of EPSC/IPSC amplitude ratios in the detailed model

For comparison with experimental data, we examine what EPSC/IPSC amplitude ratios exist for cells in the detailed model. We choose 15 cells of each type from the full-scale model (***Bezaire et al., 2016b***). These types are PYR cells and fast-firing PV+ cell types - BCs, BiCs and AACs. In doing this examination it is important to note that experimental estimates of these ratios as derived from voltage clamp recordings are not precise as there are associated experimental limitations such as due to space clamp. However, the experimental data shows that EPSCs received by PV+ cells are much larger in amplitude than EPSCs received by PYR cells, and since IPSCs received by PV+ and PYR cells are similar in amplitude, the experimental limitations are moot as it is clearly the case that the EPSC/IPSC amplitude ratios for PYR cells are much less than for PV+ cells (***Huh et al., 2016***).

In considering the detailed model, several aspects need to be taken into consideration. First, in the detailed model, we consider fast-firing PV+ cell types as BCs, BiCs or AACs in different combinations (see main text). Next, with the detailed model, morphological representations of cells are used and there are eight different inhibitory cell types. These different inhibitory cell types synapse onto different parts of the PYR cell tree and as such, IPSCs onto PYR cells would be attenuated by different amounts when examining synaptic currents at their somata. We note that to directly compare synaptic currents from the experiments with the detailed model, one could consider performing a voltage clamp on model cells and separately examining EPSCs and IPSCs as done experimentally, but one would additionally need to separate IPSCs that are due to the different inhibitory cell types to consider PV+ or PYR cells. Undertaking this in the detailed model would be a highly non-trivial endeavour, and indeed, decades of research has uncovered the richness and complexities of dendritic integration (***Stuart and Spruston, 2015***). Thus, since we know that the EPSC/IPSC amplitude ratios are very different on PYR and PV+ cells, we focus on EPSCs and IPSCs on either PYR or PV+ cells at somatic locations without trying to compensate for voltage clamp or attenuation issues due to different synaptic input locations from the different cell types. From the consideration that the comparison is with experiment, we consider that EPSCs onto the different cell types are due to inputs from PYR cells and EC and CA3, whereas IPSCs are from the various inhibitory cell types of the detailed network model (***Bezaire et al., 2016b***). As we are mainly considering comparisons with the minimal model, we consider IPSCs that are due to PV+ fast-firing cell type could encompass BCs, BiCs and AACs.

The network clamp tool in SimTracker enables extraction of a particular cell from the full-scale model while keeping synaptic properties (***Bezaire et al., 2016a***). We network clamp each of the 15 selected cells of each type for 1000 msec and detect the peak EPSCs and IPSCs by implementing the minimum peak distance algorithm in MATLAB. For EPSC/IPSC amplitude ratio calculations for a specific cell, all excitatory currents are summed and divided by the summed inhibitory currents that the cell receives. For EPSC/IPSC amplitude ratios on to PYR cells, IPSCs due to only BCs, only BiCs, a combination of BCs and BiCs, a combination of BCs/BiCs/AACs, and all inhibitory cells are shown in ***Table 3***. We note that there is no EPSC/IPSC amplitude ratio consideration of AACs to themselves as there are no AAC to AAC synapses in the detailed model. When there is a combination, the ratio calculations are based on dividing the mean EPSCs by mean IPSCs, after summing IPSCs from each PV+ cell type. The EPSCs are flipped before peak detection for its mechanistic advantage using the MATLAB code. All 225 (15×15) combinations of EPSC/IPSC amplitude ratios in each BC/BiC/PYR and BC/AAC/PYR populations as well as 3375 (15×15×15) combinations in BC/BiC/AAC/PYR are examined, and they are in accordance with the experimental data. The mean EPSC/IPSC amplitude ratios and their standard deviations for the various cell types are given in ***Table 3***. Voltage recordings and currents plots from the 15 chosen cells can be accessed at https://osf.io/yrkfv/. The scripts for the EPSC/IPSC amplitude ratio calculations can be found at https://github.com/FKSkinnerLab/CA1_SimpleDetailed.

#### Parametric explorations in the segment model

To generate the heatmaps of ***Figure 9*** we use the following process on the created segment model. We perform exhaustive parametric explorations of the theta power dependence on the excitatory drives in the segment model. We vary the EC/CA3 to PYR cell synaptic conductance *g*_*ec*/*ca*3–*pyr*_, the PYR-PYR synaptic conductance *g_pyr–pyr_* and the level of external stimulation, which represents the firing rate of our external EC and CA3 cells. For every pair of *g_pyr–pyr_* and *g*_*ec*/*ca*3–*pyr*_, we search for the level of external stimulation that maximizes the normalized theta power. The normalized theta power is defined as the maximum theta power (net theta power) in the power spectrum, divided by the mean power across all frequencies. We search a range of 0.15-0.65 Hz of stimulation per network (below that range the network is inactive, above that range the network is hyper-active). We plot the value of that maximum normalized theta power in ***Figure 9***Bi, and the corresponding stimulation required to reach that value in ***Figure 9***Biii. Every pair of *g_pyr–pyr_* and *g*_*ec*/*ca*3–*pyr*_ corresponds to a specific conndata#.dat file. These conndata#.dat files should be created and stored under the “datasets” directory of the CA1 repository. The code for the generation of the heatmaps of ***Figure 9***B can be found here: https://github.com/alexandrapierri/CA1-Segment-Microcircuit

#### Current extractions and linear regression in the segment model

As described above for ratio calculations in the detailed model, we use the network clamp tool of SimTracker to extract PSCs delivered to the PYR cells in the model from all other cells in the network and the external drives. We examine the PSCs received by 10 PYR cells for each of the 50 networks underpinning the heatmaps of ***Figure 9***B. we calculate the mean current amplitude for each of the 10 cell over a 4sec simulation period, and refer to this as the net current. We take the average and standard deviation of the net current of the 10 cells and plot it against the frequency of that network (***Figure 10***C).

As we examine 10 cells per network and we have 50 networks, this gives as a total of 500 network clamp simulations which corresponds to analysis of 500 cells’ input currents. To perform a linear regression of net current vs network frequency, we use custom MATLAB code which can be found here: https://github.com/alexandrapierri/CA1-Segment-Microcircuit. The correlation coefficient between theta frequency and net current (*ρ*) and the p-value for testing the hypothesis of no correlation (null hypothesis) against the alternative hypothesis of a nonzero correlation, are estimated using MATLAB’s built-in functions.

#### Power analysis and signal filtering

To analyze the signal power we used the Welch’s Periodogram, a method for estimating power spectra based on FFT analysis https://ccrma.stanford.edu/~jos/sasp/Welch_s_Method.html. To filter the LFP signal for theta we used a broadband filter with stopband frequencies ±1 Hz and passband frequencies ± 1.75 Hz from the peak theta frequency as derived from the Welch’s Periodogram.

#### High performance computing simulations

We implement our simulations on Scinet (***Loken et al., 2010***; ***Ponce et al., 2019***) on the Niagara clusters, using 10-12 nodes per simulation with 40 cores per node. Each network simulation takes approximately 8 hours real time to be executed. The results we present in this study are the distillation of approximately 300 network simulations requiring a total of 150 core years processing power on the clusters.

## Acknowledgements

FKS and APC thank Ivan Soltesz for feedback on early versions of this work. APC thanks Ivan Raikov for help with code debugging of the detailed model, and FKS thanks Sylvain Williams for feedback comments on the paper.

Computations were performed on the Niagara supercomputer at the SciNet HPC Consortium. SciNet is funded by: the Canada Foundation for Innovation; the Government of Ontario; Ontario Research Fund - Research Excellence; and the University of Toronto.

---

**Figure 3–Figure supplement 1.**
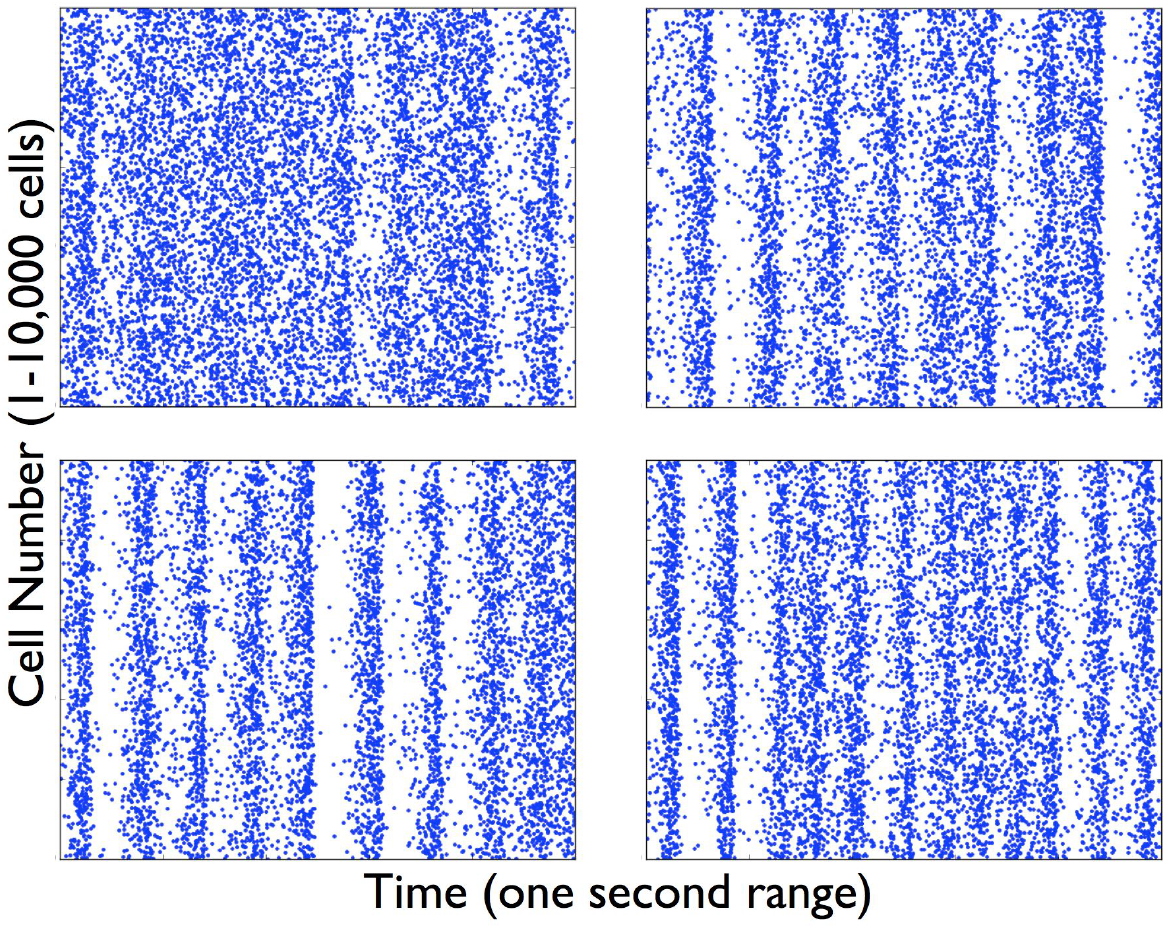
Loss of Rhythm - Raster plots of PYR cells in heterogeneous E-I networks. Simulations of E-I networks with 10,000 heterogeneous PYR cells and 500 PV+ cells produce PYR cell raster plots as shown here with a one second time range. The specific examples are labelled as (*R-supp*) in **Table 2** and refer to the following sets: MLH (top-left), HMH (top-right), MLM (bottom-left), LLH (bottom-right).

**Figure 6–Figure supplement 1.**
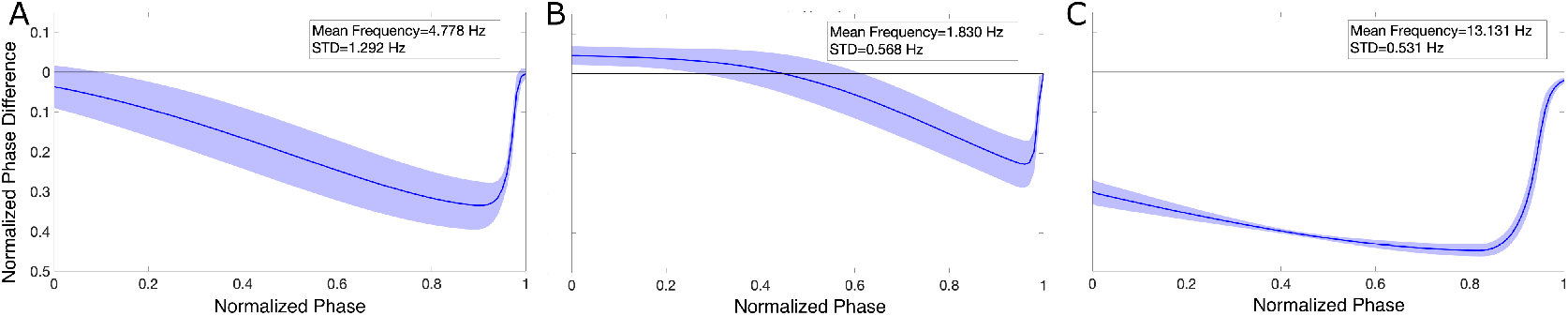
PRCs calculated with a 20 pA input show similar features in the three PYR cell populations. Mean and standard deviation of the PRCs calculated for PYR cell models from each of the three heterogeneous E-I network cases (*MMH* in panel **A**, *HML* in panel **B**, and *LML* in panel **C**) with an input current of 20 pA show similar patterns to those seen in Figure 6. Insets include mean and standard deviation of the individual firing frequencies of the PYR cells.

**Figure 8–Figure supplement 1.**
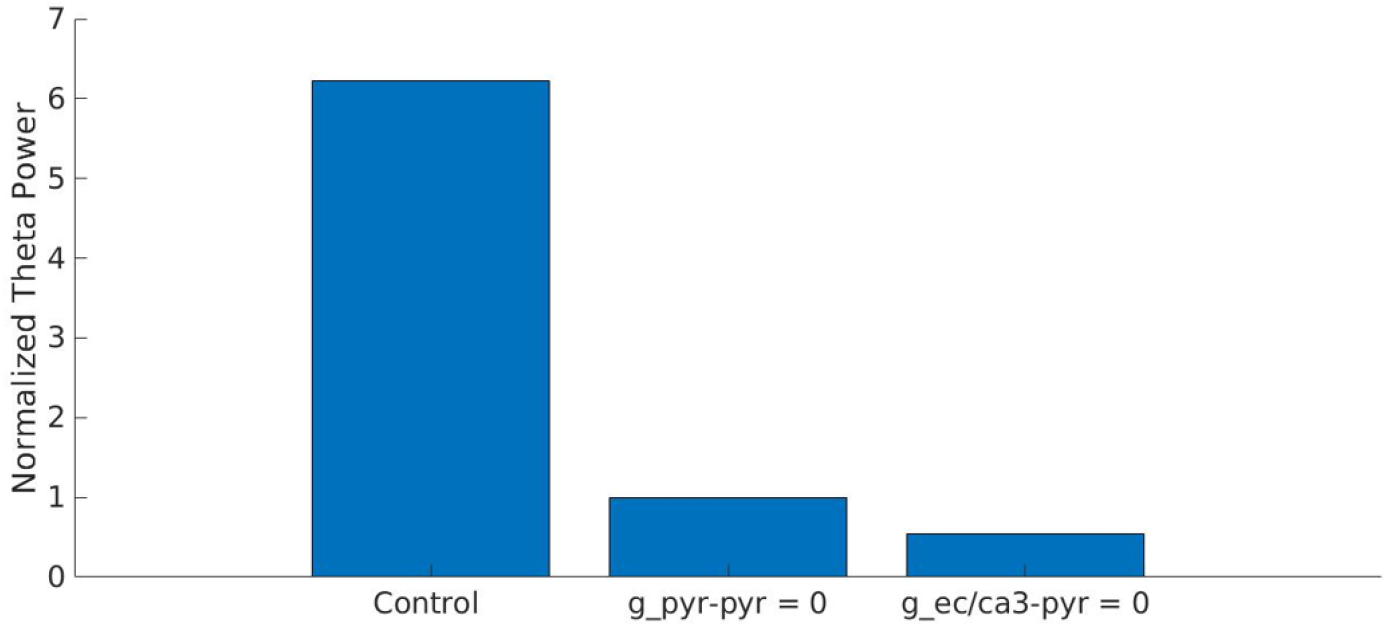
Recurrent excitation and feed-forward external drive to the PYR cells is needed for theta rhythms. Normalized theta power of the segment model (**Figure 8**A) with parameter values as shown in **Figure 8**B is eliminated with the removal of feed-forward external drive and recurrent excitation to the PYR cells, i.e., *g_pyr–pyr_* and *g*_*ec*/*ca*3–*pyr*_ set to zero.

**Figure 9–Figure supplement 1.**
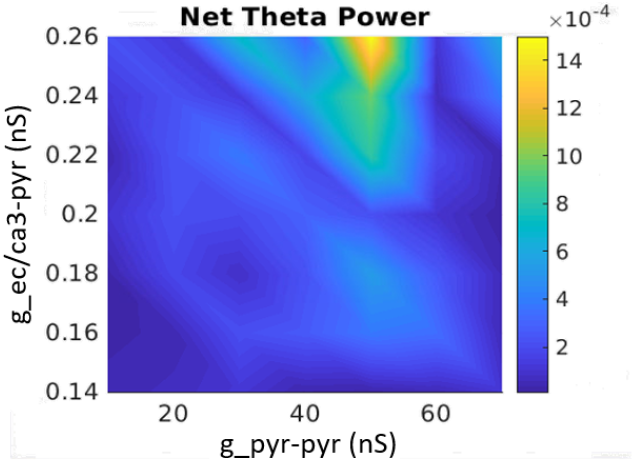
Dependence of net theta power on the PYR cells’ excitatory drives. Heatmaps of net theta power as a function of *g_pyr–pyr_* and *g*_*ec*/*ca*3–*pyr*_.

**Figure 9–Figure supplement 2.**
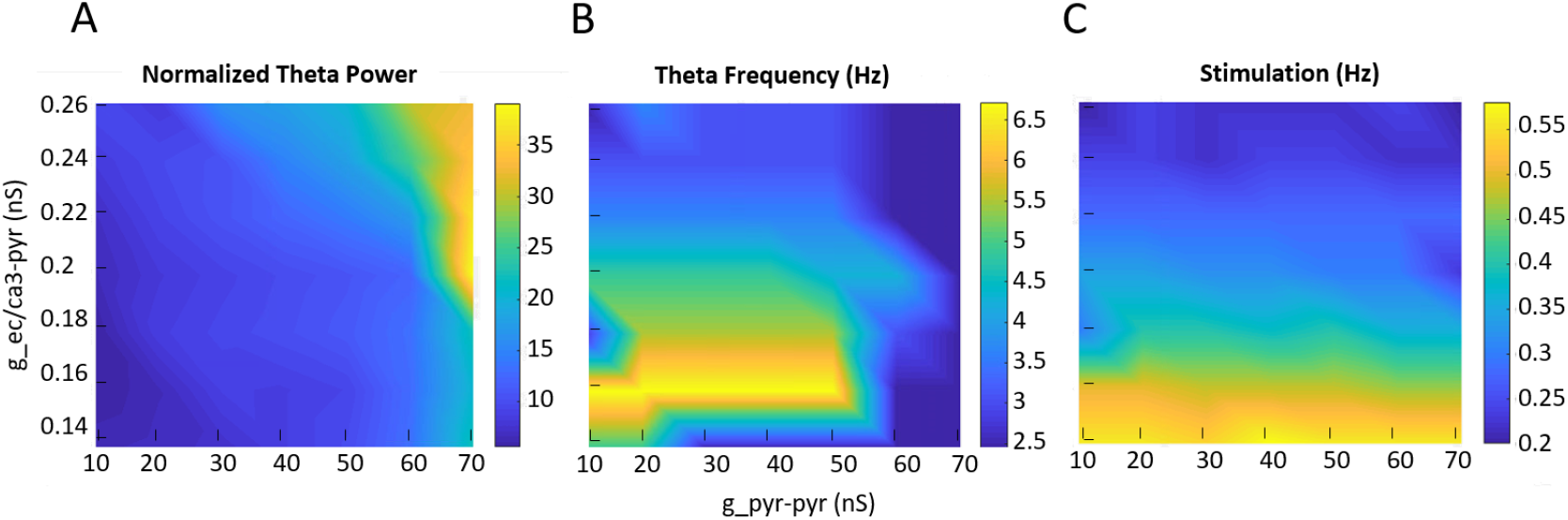
Dependence of theta and delta power on the PYR cells’ excitatory drives. Heatmaps of Normalized theta and delta power, frequency and afferent input stimulation as a function of *g_pyr–pyr_* and *g*_*ec*/*ca*3–*pyr*_.

**Figure 9–Figure supplement 3.**
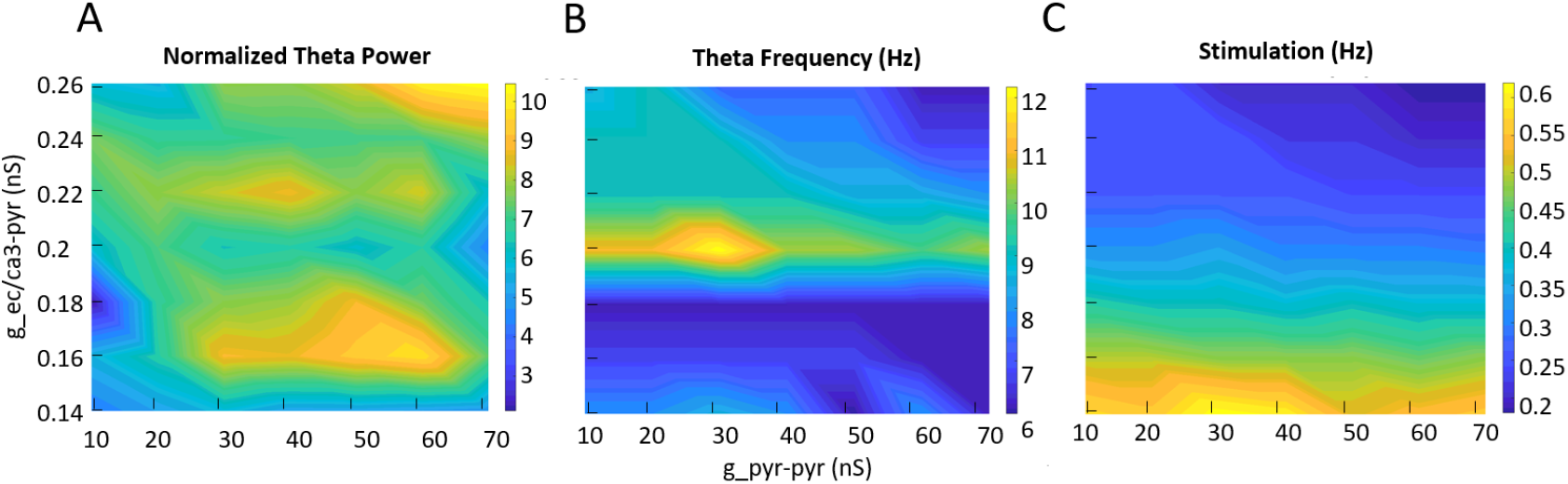
Dependence of “high” theta (6-12Hz) power on the PYR cells’ excitatory drives. Heatmaps of Normalized “high” theta power, frequency and afferent input stimulation as a function of *g_pyr–pyr_* and *g*_*ec*/*ca*3–*pyr*_.

